# Diversifying the Northern Neotropics: Phylogenomics and Evolutionary History of the Early-Diverging Herichthyine Cichlids *Thorichthys* and *Trichromis*

**DOI:** 10.64898/2026.06.05.730467

**Authors:** Diego J. Elías, Fernando Alda, Isaí Betancourt-Resendes, Alejandro Díaz-Flores, Omar Domínguez-Domínguez, Sheila Rodríguez-Machado, Ernesto Velásquez-Velásquez, Kyle R. Piller, Wilfredo A. Matamoros, Susan F. Mochel, Kevin A. Swagel, Prosanta Chakrabarty, Caleb D. McMahan

## Abstract

Among Neotropical cichlids the tribe Heroini exhibits substantial ecological diversity and is one of the dominant fish groups across northern Neotropical riverscapes. The majority of studies on heroine cichlids have focused on macroevolutionary patterns but the role of geological and ecological factors shaping lineage diversification within the tribe remains poorly understood. Here we used ultraconserved elements (UCEs) to infer a taxonomically complete and geographically comprehensive phylogenomic framework of the early-diverging herichthyine sister genera *Thorichthys* and *Trichromis* and to comparatively investigate their evolutionary and biogeographic histories. All our phylogenomic hypotheses support the monophyly of both genera and two species subgroups within *Thorichthys*. Our results provide evidence of a) unrecognized diversity within *Trichromis salvini*, b) uncertainty in species boundaries in *Thorichthys*, and c) the first report of ghost introgression in fishes of the northern Neotropics. Additionally, our results demonstrate the importance of the Papaloapan and Coatzacoalcos watersheds for fish diversification in the region. Finally, we show patterns consistent with ecological divergence during the evolution of this group, particularly among sympatric species. Altogether, these patterns suggest that lineage diversification in northern Neotropical cichlids has been driven by the interaction of geological restructuring, climatic and sea-level oscillations, and heterogeneous ecological pressures.

## INTRODUCTION

In aquatic ecosystems, ecological radiations are well documented in lacustrine environments, where colonization-associated ecological opportunity can trigger rapid ecological divergence (e.g., tropical cichlids, sticklebacks, and Arctic char; McGee, Schluter & Wainwright, 2013; Seehausen, 2015; Doenz et al., 2019; Torres-Dowdall & Meyer, 2021; Wagner, 2021). In contrast, riverine systems traditionally have been seen as less conducive to ecological diversification due to their limited potential for ecological opportunity (Seehausen, 2015). However, recent studies have challenged this view by providing examples of ecological diversification in riverine fishes (e.g., Burress et al., 2018; Levin et al., 2021). In riverine environments, allopatric speciation driven by geological processes is considered the primary mechanism of diversification (Albert et al., 2020; Walters et al., 2026). Among Neotropical freshwater fishes, cichlid diversity is dominated by members of the tribes Geophagini, Cichlasomatini, and Heroini, with Heroini being the only tribe to have colonized the northern Neotropics (Smith, Chakrabarty & Sparks, 2008; McMahan et al., 2015; López-Fernández 2021). Diversification in Heroini unfolded mainly in two hydrological systems, the Usumacinta and San Juan watersheds, located within the Maya and Chortis blocks, respectively (Říčan et al., 2013; McMahan et al., 2015; Říčan et al., 2016; Tagliacollo et al., 2017). In northern Neotropical heroine cichlids, present-day ecomorphological diversity has been hypothesized to arise through ecological release experienced by ancestral lineages after colonization of the region (Říčan et al., 2013, 2016; Burress & Tan, 2017; Arbour et al., 2020; López-Fernández, 2021).

Because evolutionary studies of Heroini have largely focused on macroevolutionary patterns (e.g., Arbour et al., 2020; Mejía et al., 2022; Nicholas & López-Fernández, 2024), our understanding of the drivers of *in situ* diversification remains limited. Only a few heroine genera have been investigated in detail, and evidence suggest that both ecological (sympatric) and geological (allopatric) processes may have contributed to their diversification (e.g., Musilová et al., 2015; Pérez-Miranda et al., 2020a, b; Torres-Dowdall & Meyer, 2021; Leal-Cardín et al., 2024; Hulsey et al., 2026).

The herichthyines are a nested radiation within Heroini, and are hypothesized to have diversified within the Usumacinta’s evolutionary center located in the Maya block (e.g., Říčan et al., 2016). Herichthyines is comprised of 52 species in 16 genera (McMahan et al., 2015; Říčan et al., 2016), of which *Thorichthys* is one of the most species-rich, comprising nine valid species naturally distributed across the Atlantic slope from the Papaloapan River in Mexico to the Motagua River in western Honduras with translocated populations in Mexico (e.g., north of the Trans-Mexican Volcanic Belt TMVB; Fig. 1, Fig. S1; McMahan et al., 2015; Artigas Azas, 2021). Although most species are allopatrically distributed, syntopic occurrences have been documented, including *T*. *callolepis* and *T*. *panchovillai* in the upper Coatzacoalcos, and *T*. *helleri*, *T*. *pasionis* and *T*. *meeki* in the Usumacinta watershed (Del Moral Flores et al., 2017; Elías et al., 2025; Figs. S1, S2).

**Fig. 1.**
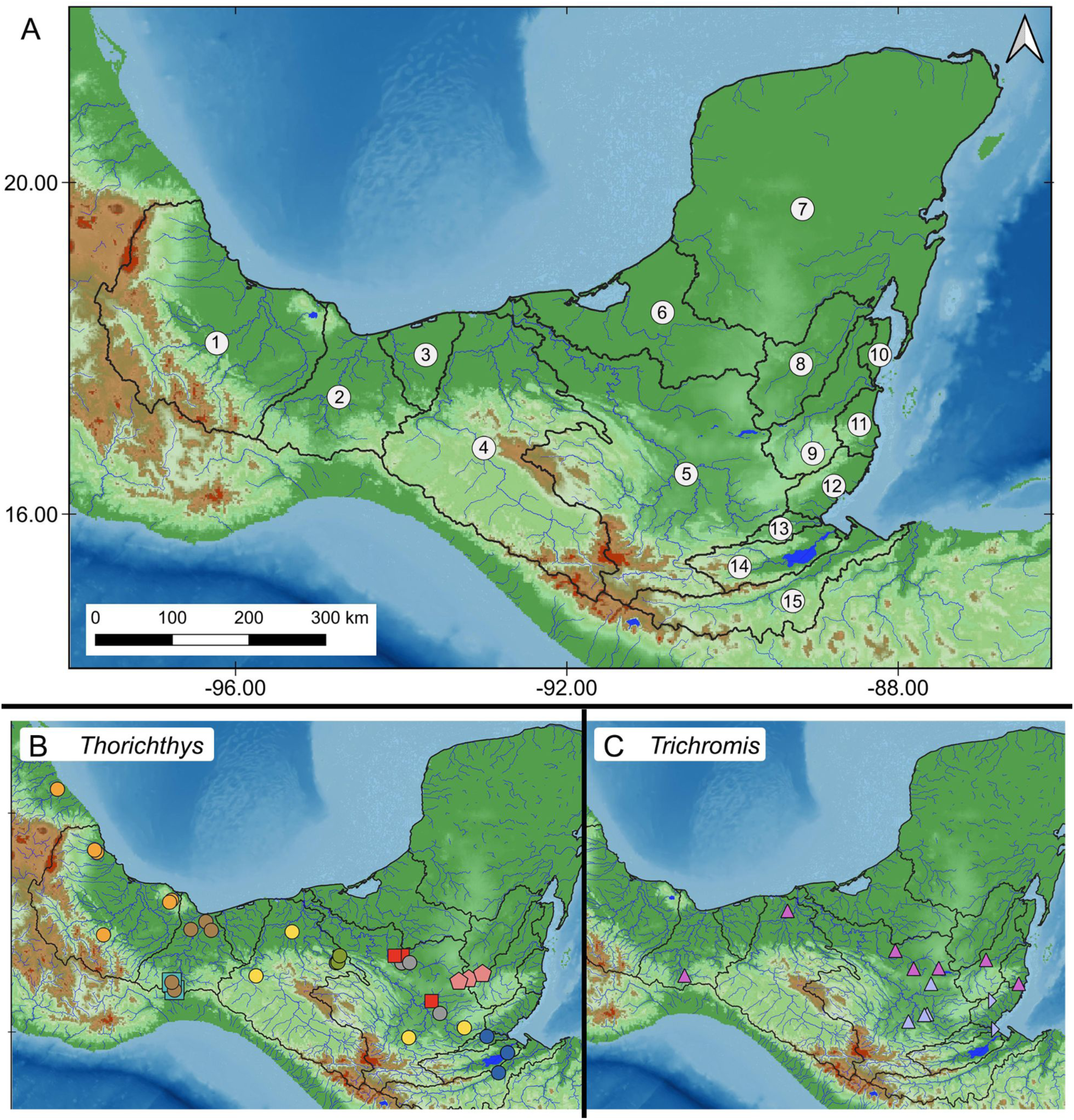
A) Map of the northern Neotropical region showing the major watersheds (black lines and numbers) where the early-diverging herichthyines are distributed: 1) Papaloapan, 2) Coatzacoalcos, 3) Tonalá, 4) Grijalva, 5) Usumacinta, 6) Candelaria-Laguna de Términos, 7) Yucatan Peninsula, 8) Hondo, 9) Mopán, 10) Northern Belize, 11) Central Belize, 12) Southern Belize, 13) Sarstún, 14) Polochic-Cahabón-Lago de Izabal, and 15) Motagua. B-C) Geographic location of the samples used for phylogenomic analyses (symbol shape, color, and watershed code in parentheses): B) *Thorichthys affinis* (salmon pink pentagons: 5 & 9), *T*. *aureus* (blue circles: 13-15), *T*. *callolepis* (teal squares: 2), *T*. *maculipinnis* (orange circles: 1), *T*. *meeki* (red squares: 5), *T*. *helleri* (yellow circles: 4, 5), *T*. *panchovillai* (brown circles: 2), *T*. *pasionis* (gray circles: 5), *T*. *socolofi* (green circles, 4). C) *Trichromis salvini* (magenta upright triangles: 2, 4, 5, 9, 12; light violet upright triangles: 5; light violet right-pointing triangles: 12, 14)

*Thorichthys* species are well known for their vibrant coloration patterns and have been popular among aquarium hobbyists for over a century (Brind, 1918; Artigas Azas, 2008). For example, the Firemouth Cichlid, *T*. *meeki*, possesses a bright red coloration on the lower portion of the jaw that is displayed during breeding or protecting territory (Konings, 1989). The genus is diagnosed by several morphological characters, including scale-less dorsal and anal fins, five mandibular pores (vs. four in all other herichthyine genera), and typically 12 precaudal vertebrae (Miller & Taylor, 1984; McMahan et al., 2015). Phylogenetic and morphological evidence supports the recognition of two clades within *Thorichthys*, the “helleri” and “meeki” species subgroups. The “helleri” subgroup was coined by Miller and Taylor (1984) and includes six species: *T*. *helleri*, *T*. *maculipinnis*, *T*. *callolepis*, *T*. *panchovillai*, *T*. *socolofi*, and *T*. *aureus*. The same authors identified a related cluster of *Thorichthys* species, including *T*. *meeki*, *T*. *pasionis*, and *T*. *affinis*. The unpublished thesis of López Segovia (2021) referred to this clade as the “meeki” subgroup. The two species subgroups differ on their gill raker and anal-fin spine counts, lower jaw and pectoral fin lengths, and lower pharyngeal jaw morphology (Miller & Taylor, 1984). Phylogenetic analyses with varying taxonomic coverage have consistently recovered reciprocal monophyly among these species subgroups; however, the inferred relationships among species within each clade have varied across studies (López-Fernández et al., 2010; McMahan et al., 2015; Říčan et al., 2016; López Segovia, 2021; Elías et al., 2025).

In contrast to the species-rich genus *Thorichthys*, its sister genus *Trichromis* is monotypic, containing only *T*. *salvini*. Despite its lack of diversification, *T*. *salvini* shares a broadly overlapping distribution with *Thorichthys*, except that *T*. *salvini* does not extend into the Motagua River (Fig. S3), and exhibits similarly vibrant coloration patterns characterized by red, yellow, and blue hues. The pronounced asymmetry in species richness between these herichthyine genera raises intriguing questions about the processes promoting diversification in riverine ecosystems in the region.

In this study, we investigate the evolutionary history of the early-diverging herichthyine sister genera *Thorichthys-Trichromis* using a taxonomically and geographically comprehensive phylogenomic framework. We integrate biogeographic and ecomorphological analyses to evaluate the relative roles of allopatric and ecological processes in shaping diversification in northern Neotropical cichlids.

## MATERIAL AND METHODS

### Taxon Sampling

Our taxon sampling included a total of 11 species of northern Neotropical cichlids comprising all valid species of the herichthyine genera *Thorichthys* (nine species), *Trichromis* (one species), and *Chiapaheros* (one species), with the latter selected as an outgroup based on previous phylogenomic hypotheses (see Ilves et al., 2018; Alda et al., 2021; Table S1). The molecular data set was comprised of a total of 54 individuals (i.e., tissue samples) representing 40 samples from all valid species of the genus *Thorichthys*: *T*. *affinis* (n =3), *T*. *aureus* (n =3), *T*. *callolepis* (n =3), *T*. *helleri* (n =5), *T*. *maculipinnis* (n =8), *T*. *meeki* (n =4), *T*. *panchovillai* (n = 8), *T*. *pasionis* (n =3), *T*. *socolofi* (n =3), and 13 samples of the monotypic *Trichromis salvini* from across its distribution (Fig. 1B, C). Additionally, we included one sample of *Chiapaheros grammodes*. Samples were provided by ichthyological collections of the Louisiana State University Museum of Natural Science (LSUMZ), Southeastern Louisiana University Vertebrate Museum (SLU), Field Museum of Natural History (FMNH), and the Museo de Zoología Universidad de Ciencias y Artes de Chiapas, Mexico (MZ-UNICACH). Acronyms follow Sabaj (2020).

### DNA Extraction and amplification of cytochrome b

We extracted genomic DNA using the DNeasy Blood & Tissue kit (QIAGEN) following the manufacturer’s protocol. The quality of our extractions was visually assessed using a 2% agarose gel, and the DNA concentration was quantified using a Qubit® 2.0 fluorometer (Invitrogen). To corroborate taxonomic identity of our samples, we amplified and sequenced the mitochondrial cytochrome b gene (Cytb) following the methods described in Elías et al. (2025), and we compared the sequences with the NCBI database using the BLAST algorithm online (https://blast.ncbi.nlm.nih.gov/Blast.cgi).

### UCE Library Preparation

To generate a dataset of ultraconserved elements (UCEs; Faircloth et al., 2012; https://www.ultraconserved.org), we used 300 ng of DNA per sample as starting material. We sheared the DNA to a target fragment size between 400 – 600 base pairs (Faircloth & Glenn, 2012) using an EpiSonic Multi-functional bioprocessor (EpicGenTek). After DNA was fragmented, we used the Kapa Hyper Prep Kit (Kapa Biosystems) to construct dual index libraries, which were pooled and enriched following the protocols described in https://www.ultraconserved.org/#protocols with the adjustments of Burress et al. (2018). We then used the myBaits UCE Acanthomorph 1Kv1 kit (Daicel Arbor Biosciences) to target the capture of 1,314 UCE loci across the genomes of our samples (McGee et al., 2016). Libraries were hybridized and enriched in pools of eight samples. Quality control of the final libraries was performed using Bioanalyzer (Agilent) at the LSU Genomics Core (https://genomics.lsu.edu/) prior to sequencing.

Finally, all libraries were equimolarly combined into a final pool with a concentration of 10 µM that was then sequenced on a partial lane (150 GB of raw data) in an Illumina NovaSeq X Plus Sequencing System (PE150) by Novogene (https://en.novogene.com/). Raw reads were archived in the National Center for Biotechnology Information (NCBI) sequence repository archive (SRA) under BioProject XXXXX.

### Bioinformatics

We used the software fastp to assess the quality of our demultiplexed raw reads and to remove adapter contamination and low-quality reads (Chen, 2023, 2025). After quality control, we used the PHYLUCE v1.7-1 software (Faircloth, 2016) to assemble the clean reads for each individual sample into contigs with SPAdes (Bankevich et al., 2012). Next, to identify and extract UCE loci we ran the assembled contigs for all the samples against the Acanthomorph UCE probe set, and we generated a list of all loci recovered that we used to create monolithic FASTA files matching all loci in the list. We exploded each monolithic FASTA file by locus and aligned their sequences using the software MAFFT v. 7 (Katoh & Standley, 2013) with edge trimming. Finally, we filtered the aligned loci to generate a data matrix of 75% completeness, retaining only loci present in at least 40 of the 54 samples. Summary statistics of final alignments (see Table S2) were calculated using SEGUL (Handika & Esselstyn, 2024). This dataset was used for all downstream phylogenomic analyses (concatenation and coalescent approaches, see below). Data processing steps described above follow standard protocols (see https://phyluce.readthedocs.io/) and were performed on the Grainger Bioinformatics Center Server ‘Phoebe’ at the Field Museum of Natural History.

### Phylogenomic Inference

To investigate evolutionary relationships between species and within populations of the early-diverging herichthyine genera *Thorichthys* and *Trichromis*, we inferred phylogenomic hypotheses based on concatenation and summary coalescent approaches, using IQ-TREE 3 (Wong et al., 2025) and the Accurate Species Tree EstimatoR (ASTER) package (Zhang, Nielsen & Mirarab, 2025a), respectively.

### Concatenation inference

For the concatenation approach we first performed model selection for each UCE locus recovered in our 75% complete data matrix using ModelFinder (Kalyaanamoorthy et al., 2017) implemented in IQ-TREE 3, followed by Maximum Likelihood (ML) tree inference in the same run using the option *-m TEST*. To assess node support of the inferred concatenated topology we used the Ultrafast bootstrap (UFBoot2; Minh et al., 2013; Hoang et al., 2017) with 1000 replicates each using the option *-bb 1000*.

### Species tree inference

To infer the multispecies coalescence-based species tree of *Thorichthys* and *Trichromis* we leveraged four programs available in the ASTER package that can be divided into: summary coalescent and alignment-based approaches.

#### Summary coalescent species tree

We used the programs ASTRAL-IV (Zhang, Nielsen & Mirarab, 2025a) and weighted-Astral (wASTRAL v1.23.3.7; Zhang & Mirarab, 2022), which utilize unrooted gene trees as input to estimate a species tree that is statistically consistent with the multispecies coalescent (MSC) model, assuming that gene tree discordance arises from incomplete lineage sorting (Mirarab et al., 2014; Zhang & Mirarab, 2022). These two softwares differ in how they try to minimize the effects of gene tree estimation error (Simmons & Gatesy, 2021) when inferring the species tree. In ASTRAL-IV, collapsing branches with low support (e.g., ≤ 10% support value) has been recommended (Mirarab 2023; Zhang, Nielsen & Mirarab, 2025a). This strategy has been widely used when reconstructing species trees from UCE data, often using different cutoff thresholds of support values to collapse branches (e.g., ranging from 0 to 50; Alda et al., 2021; Elías et al., 2023; Rodriguez-Machado et al., 2024; Alda et al., 2025; Liyandja et al., 2025; Borowiec et al., 2025; López-Estrada et al., 2026). Alternatively, wASTRAL uses a weighting scheme based on gene tree branch lengths and support values, avoiding the need to set a support threshold for collapsing low-support branches. This method has been shown to improve accuracy when inferring the species tree under summary coalescent approaches (Zhang & Mirarab, 2022). Additionally, these two methods differ in their estimation of branch lengths; ASTRAL-IV estimates internal and terminal branch lengths in substitutions-per-site (Zhang, Nielsen & Mirarab, 2025a), whereas wASTRAL does not estimate branch lengths for the inferred species tree (Zhang & Mirarab, 2022).

We estimated gene trees with support values (UFBoot2; 1000 replicates) for each UCE locus in the 75% complete data matrix using IQ-TREE 3 setting options *-s* and *-bb* and used them as input for both ASTRAL-IV and wASTRAL. To minimize the effect of gene tree estimation error when estimating the species tree in ASTRAL-IV, we collapsed branches with support values (UFBoot2) ≤ 10 in all estimated gene trees following Alda et al. 2021. In contrast, for wASTRAL we used the best-scoring gene trees without collapsing branches.

For both methods, a species tree was first inferred with each sample represented as a terminal tip (i.e., multiple individuals per species/lineage). This initial tree was used to identify the most inclusive sets of individuals forming strongly supported clades that may correspond to species or independent lineages within *Thorichthys* and *Trichromis*. Subsequently, we inferred a second species tree in which individuals within each of these highly supported clades were constrained to the same species or lineage.

To this end we generated a mapping file that we provided as input when running ASTRAL-IV and wASTRAL using the *-a* option. Branch support for the resulting species trees was assessed using local posterior probabilities (LPP; Sayyari & Mirarab, 2016). Additionally, we used normalized quartet scores (NQS; Sayyari and Mirarab, 2016) to investigate gene tree-species tree discordance. We annotated the species tree using the *-u 3* option in ASTRAL-IV and wASTRAL to calculate the normalized quartet scores of the main topology (i.e., the preferred species tree), its internal branches, and the two alternative quartet topologies.

#### Alignment-based species tree

We used a novel approach to species tree inference based on site patterns consistent with the incomplete lineage sorting model: the Coalescent-aware Alignment-based Species Tree EstimatoR (CASTER; Zhang, Nielsen & Mirarab, 2025b). Unlike summary coalescent methods, CASTER does not require prior gene tree estimation to estimate the species tree, as it operates directly on genome-wide sequence alignments. We implemented the two approaches available in the CASTER algorithm: CASTER-site and CASTER-pair. The first approach optimizes site patterns across quartets of species, whereas the second optimizes pairs of sites across quartets of species. CASTER-site is statistically consistent under the MSC+F84 model, allowing for changes in substitution rates in sites and branches across the species tree. In contrast, CASTER-pair is consistent with the MSC + Lumpable Markovian models (LM), in which paired sites evolve under the same substitution model but may have distinct mutation rates (see Zhang, Nielsen & Mirarab, 2025b). Branch support for the species trees was assessed using local bootstrap support (LBS) where values > 95 are considered strong support.

### Timeline of diversification of the Thorichthys-Trichromis clade

To better understand the diversification of early-diverging herichthyines, we generated a time-calibrated tree (i.e., chronogram) using a penalized likelihood approach (Sanderson, 2002; Kim & Sanderson, 2008; Paradis, 2013) implemented with the *‘chronos’* function in the R package APE (Paradis et al., 2008; Paradis & Schliep, 2019). For this analysis, we used the multiple individual species tree with branch lengths inferred in ASTRAL-IV and applied three secondary calibration points to estimate divergence times in ‘*chronos*.’

The minimum and maximum age estimates used for the three secondary-calibrations were obtained from a family level UCE topology (Elías et al. *in prep*) that was dated using four fossil calibrations points 1) Pseudocrenilabrinae less *Heterochromis* (max = 94 mya, min = 48 mya), 2) *Geophagus* + *Gymnogephagus* (max = 95 mya, min = 40 mya), 3) Cichlasomatini + Heroini (max = 94 mya, min = 40 mya), and 4) *Nandopsis woodringi* (max =46 mya, min = 3.6 mya) (McMahan et al., 2013; https://fishtreeoflife.org/fossils/nandopsis-woodringi/). Age estimates were assigned to the following nodes: 1) the most recent common ancestor (MRCA) of *Chiapaheros*-*Thorichthys*-*Trichromis*, 2) the MRCA of *Thorichthys*-*Trichromis*, 3) the MRCA of *Thorichthys* (see Figs. 2A, 3B). These nodes were selected based on previously published phylogenetic relationships across different taxonomic-scales (see Ilves et al., 2018; Alda et al., 2021; Elías et al., 2025).

**Fig. 2.**
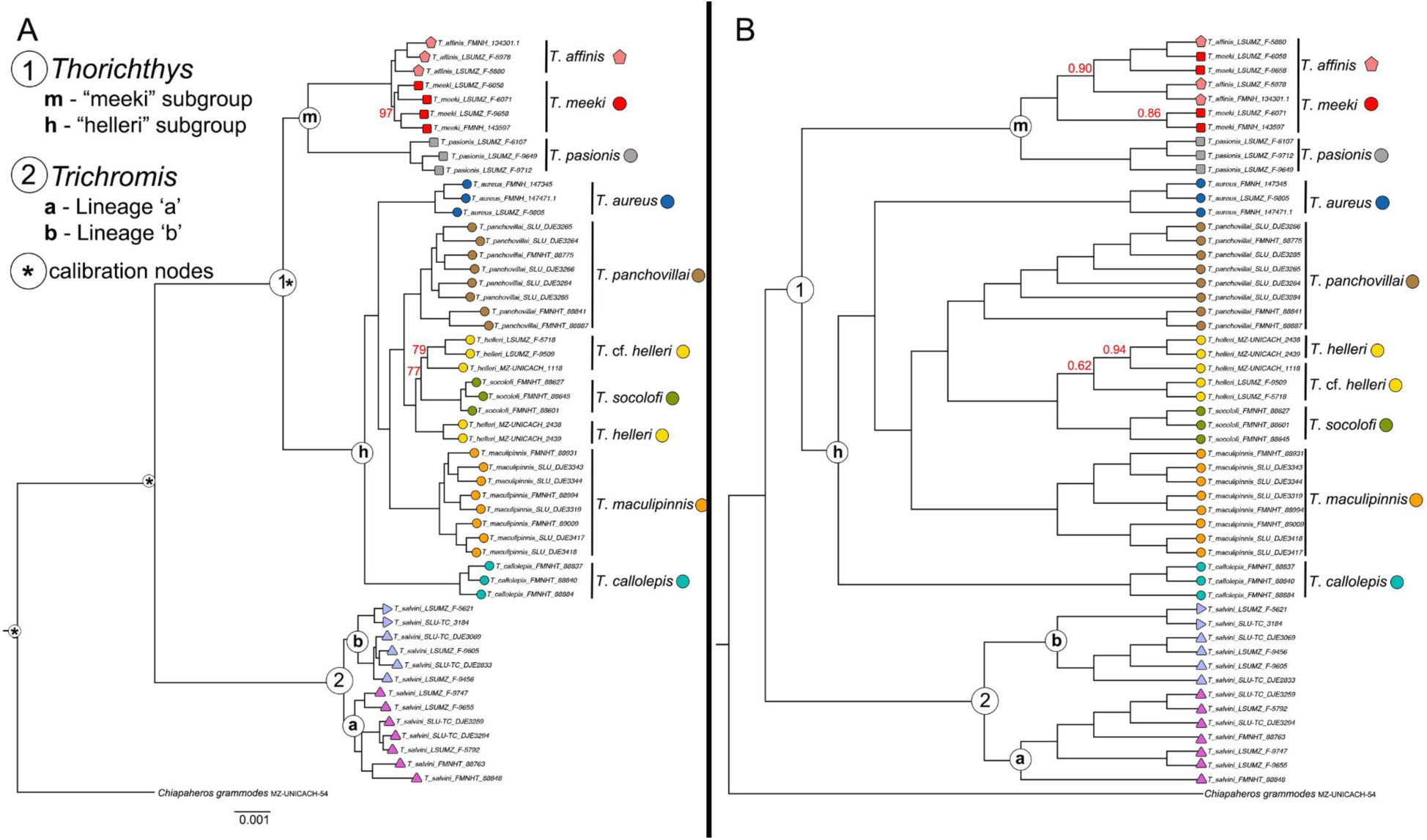
Evolutionary relationships of lineages of the early diverging Herichthyines clade *Thorichthys*-*Trichromis* (TT) inferred from the 75% complete UCE data matrix (997 UCE loci). A) Phylogenetic relationships of the TT clade inferred under a maximum likelihood approach implemented in IQ-TREE 3. All nodes subtending to species/lineages are supported by ultrafast bootstrap (UFboot2) = 100 unless indicated in the phylogeny. * denotes nodes that where secondary calibrations were added in the dating analyses. B) Cladogram of relationships of the TT clade inferred under a summary coalescent approach weighted by gene tree uncertainty (wASTRAL) implemented in ASTER. All nodes subtending to species/lineages are supported by local posterior probabilities (LPP) = 1.0 unless indicated in the phylogeny

**Fig. 3.**
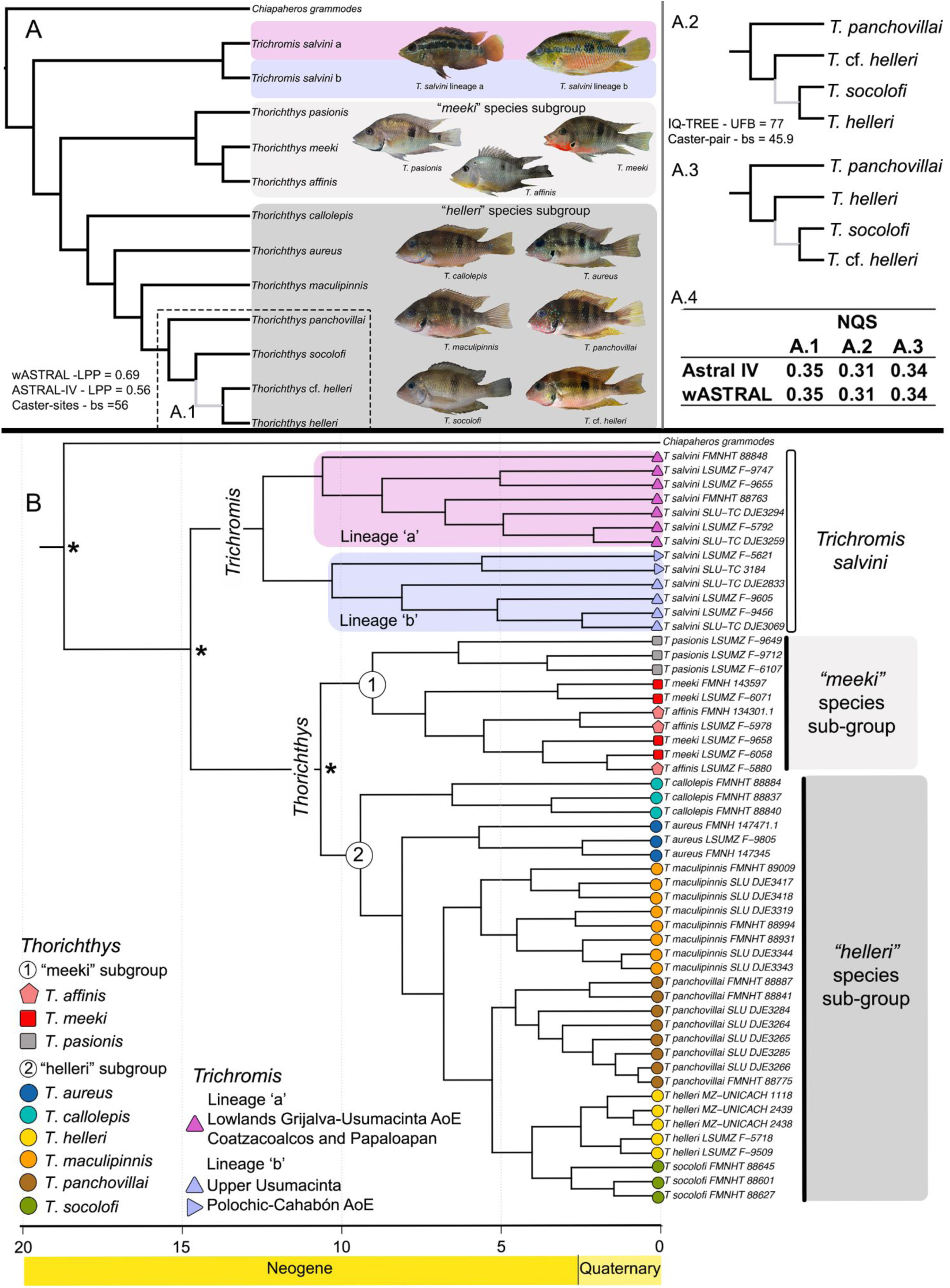
A) Species tree cladogram of the early divergent Herichthyine genera *Thorichthys* and *Trichromis*. A.1-A.3) Alternative phylogenetic relationships between *Thorichthys socolofi*, *T*. *helleri*, and *T*. cf. *helleri* and support values (UFB: IQTREE 2; LPP: ASTRAL-IV and wASTRAL; bs: Caster-pairs and Caster-sites) recovered for the contentious branch (in light grey) A.4) Normalized quartet scores (NQS) for the alternative relationships between *T. socolofi*, *T*. *helleri*, and *T*. cf. *helleri*. B) Chronogram of the early diverging herichthyines. * denotes nodes that where secondary calibrations were placed for the dating analyses

We conducted a sensibility analysis using four values of the smoothing parameter λ (0.1, 1, 10, a100), representing increasing rate variation across branches at lower λ values and decreasing rate variation at higher λ values (Sanders, 2002) under a “relaxed” model of substitution rate variation. We kept the dated topology with the lowest ΦIC score (Paradis, 2013) as the preferred chronogram.

Finally, we visualized patterns of diversification by constructing lineage-through-time (LTT) plots independently for the *Thorichthys* species subgroups and *Trichromis*, using the dated topology pruned to include one representative per species/lineage within each group. Lineage-through-time plots were generated using the *ltt* function in phytools 2.0 (Revell, 2024).

### Biogeography of the early-diverging Thorichthys-Trichromis clade

Landscape evolution, including the reconfiguration of geological blocks (e.g., Maya, Chortis) and eustatic sea-level changes, has played an important role in the evolutionary history of freshwater fishes in the northern Neotropics (Říčan et al., 2013; Bagley & Johnson, 2014; McMahan et al., 2017; Tagliacollo et al., 2017; Pérez-Miranda et al., 2020; Elías et al., 2023). Accordingly, allopatric speciation, particularly through processes such as river capture, is widely considered as the primary mode of geographic diversification in Neotropical freshwater fishes (Albert et al., 2020). However, incomplete taxonomic sampling across phylogenetic hypotheses of heroine cichlid genera have limited our ability to reconstruct the biogeographic history of this radiation.

We integrated the time-calibrated phylogenomic hypothesis with present-day species distribution derived from curated museum records obtained through the Global Biodiversity Information Facility (GBIF, 2026 a, b) to infer the biogeographic history of *Thorichthys* and *Trichromis*. To specifically explore how landscape evolution has influenced diversification, we generated paleogeographic maps for the main (i.e., Maya and Chortis blocks) and adjacent (i.e., Oaxaquia-Guerrero) geological terrains (Keppie, 2004; Marshall, 2007) where this clade is currently distributed. These maps were generated in GPlates v2.5 (Müller et al., 2018; Müller et al., 2019) following the Plate Reconstructions workflow by Sheppard, Matthews & Whittaker (https://tutorials.gplates.org/), and using the time-calibrated paleogeographic reconstructions of Cao et al. (2017). Additionally, we assessed the extent of marine transgression (i.e., inundated area) and retreat (i.e., exposed continental shelf) in this region. We generated shapefiles representing the minimum and maximum sea levels during the timeframe of diversification of the *Thorichthys-Trichromis* clade. Shapefiles were derived from global ocean and terrain models obtained from the General Bathymetric Chart of the Oceans (GEBCO, 2025), with sea-level estimates based on Miller et al. (2020) and compared with estimates from Haq, Hardenbol & Vail (1987) from the dataset published in Hill (2015).

### Ecomorphological differentiation within the Thorichthys-Trichromis clade

It has been proposed that the ecological release experienced by heroine cichlids after their colonization of Middle America boosted their diversification rates, accumulating more species than expected based on their estimated age, and occupying a broader morphospace of ecomorphological key traits (McMahan et al., 2013; Arbour & López-Fernandez, 2016; Burress & Tan, 2017; López-Fernandez, 2021). However, macroevolutionary studies of heroine cichlids have only included a few selected species of the species-rich genus *Thorichthys*, and its diversity has been characterized as ecologically conservative with all the species described as substrate-sifting cichlids (López-Fernández et al., 2014; Říčan et al., 2016; Burress & Muñoz, 2023). Nevertheless, earlier studies found interspecific differences in traits associated with diet (e.g., lower pharyngeal jaw) between the two *Thorichthys* species subgroups (Miller & Taylor, 1984). Here, we investigate whether the species-rich genus *Thorichthys* exhibits unrecognized ecomorphological diversity in traits linked to ecological function, despite its apparent ecological conservatism (i.e., substrate-sifting), by comparing all nine species across the two subgroups and with its monotypic sister genus *Trichromis* within an evolutionary-explicit framework.

We performed linear measurements for six traits: head length [HL], head height [HH], snout length [SnL], eye diameter, body depth [BD], and standard length [SL] that have been linked to ecomorphological performance in cichlid fishes (López-Fernández et al., 2014). All measurements were taken to the nearest 0.01 mm with a Mitutoyo vernier caliper. We collected linear morphometric data from a total of 227 individuals: *Thorichthys affinis* (n = 18), *T*. *aureus* (n = 22), *T*. *callolepis* (n = 18), *T*. *helleri* (n = 28), *T*. *maculipinnis* (n = 23), *T*. *meeki* (n =24), *T*. *panchovillai* (n = 19), *T*. *pasionis* n = 22), *T*. *socolofi* (n = 15), and *Trichromis salvini* (n = 38). Specimens used for the meristic measurements are archived in the ichthyological collections of the Louisiana State University Museum of Natural Science (LSUMZ), Southeastern Louisiana University Vertebrate Museum (SLU), Field Museum of Natural History (FMNH), Museo de Zoología Universidad de Ciencias y Artes de Chiapas, Mexico (MZ-UNICACH), Museum of Zoology University of Michigan (UMMZ), and the Biodiversity Center at the University of Texas at Austin (THNC) (Table S3).

Morphological analyses were performed in the statistical software R ver. 4.1.1. First, we log-transformed the six linear measurements using the *log* function and generated size-corrected models for four of the measurements using the *lm* function. Specifically, HH, SnL, and ED were size-corrected using HL, whereas BD was size-corrected with SL, as a proxy for body elongation (i.e., finesse ratio; Aguirre et al., 2016; McMahan et al., 2019). Then, we extracted the residuals from these models using the function *residuals* and used them in subsequent analyses. We performed a principal components analysis (PCA) on the residuals of HH, SnL, ED, and BD and calculated the centroids for *Trichromis salvini* and the *Thorichthys* “meeki”, and “helleri” species subgroups.

Additionally, we examined whether two traits associated with ecological function in fishes provide insight into ecological diversification within early diverging herichthyines. We measured eye diameter, linked to visual performance under different light conditions (Schmitz & Wainwright, 2011), and snout length, associated with diet (Cochran-Biederman & Winemiller, 2010). We used our dated phylogeny to generate phenograms to visualize how variation in these traits is distributed across the phylogeny. We plotted the average residual value for each trait using the *phenogram* function in phytools to evaluate whether the observed variation in eye diameter and snout length showed phylogenetic signal or independence. Finally, we measured phylogenetic signal using Pagel’s λ (Pagel, 1999) implemented in the ‘*phylosig*’ function in phytools, to test whether trait variation is phylogenetically structured or independent of the evolutionary history of the group.

## RESULTS

### Alignment summary

We assembled a total of 1084 UCE loci from quality-filtered reads across 54 samples. From these, a total of 997 UCE loci were retained in our 75% matrix, with an average length of 952.44 base pairs (bp) per locus (SD = 144.24; min = 198, max = 1447 bp). Our final alignment length was 949,579 bp with a total of 904,068 (95.21%) conserved sites and 45,511 (4.79%) variable sites (VS), with an average of 45.61 VS per locus (SD = 18.68; min = 9, max = 173). Of these, 25,613 sites (2.70%) were parsimony informative (PI), with an average of 25.69 PI sites per locus (SD = 12.48; min = 1, max = 89).

### Phylogenomic inference

Our phylogenomic inference algorithms IQTREE 3 (i.e., concatenation), ASTRAL-IV, wASTRAL (i.e., summary coalescent species tree), and CASTER-site and CASTER-pair (i.e., alignment-based species tree) largely recovered the same topological relationships between and within species of early-diverging herichthyines (Figs. 2, 3) with a few relationships with relatively low support recovered across all algorithms (see below; Figs. 2, 3A).

All analyses unambiguously recovered the reciprocal monophyly of the two genera. *Trichromis salvini* was split into two well supported lineages (UFBoot2 = 100, LPP =1.0, LBS = 100; Figs. 2, 3). One lineage, *Trichromis salvini* ‘a’, is widespread from the Papaloapan, Coatzacoalcos, and Tónala river basins in southern Mexico and across the lower reaches of the Grijalva-Usumacinta AoE *sensu* Elías et al. (2020) whereas the second lineage, *T*. *salvini* ‘b’, is restricted to the Upper Usumacinta endemic area and the Polochic-Cahabón area of endemism (AoE) *sensu* Elías et al. (2020) (Fig. S3).

All analyses recovered samples of the genus *Thorichthys* as monophyletic and unambiguously identified two reciprocally monophyletic groups with high support (UFBoot2 = 100, LPP = 1.0, LBS = 100). One clade, the “meeki” subgroup, comprised all samples of *T. affinis*, *T*. *meeki*, and *T*. *pasionis* (UFBoot2 = 100, LPP = 1.0, LBS = 100; Figs. 2, 3). Relationships inferred within the ‘meeki’ subgroup recovered *T. pasionis* as sister to a clade comprised of *T*. *affinis* and *T*. *meeki,* with high support (UFBoot2 = 100, LPP = 1.0, LBS = 100; Figs. 2, 3). Interestingly, our concatenation analysis recovered all samples of *T*. *affinis* and *T*. *meeki* as reciprocally monophyletic (Fig. 2A) in contrast to our species tree inferences (i.e., ASTRAL-IV and wASTRAL), which do not recover the monophyly of these species (Fig. 2B).

The other clade, the “helleri” subgroup, comprises all the remaining diversity of the genus, including species: *T*. *aureus*, *T*. *callolepis*, *T*. *helleri*, *T*. *maculipinnis*, *T*. *panchovillai*, and *T*. *socolofi* (UFBoot2 = 100, LPP = 1.0, LBS = 100; Figs. 2, 3). Within the ‘helleri’ species subgroup, all our analyses consistently inferred *Thorichthys callolepis* as the earliest diverging species of this clade. *Thorichthys aureus* was recovered with high support (UFBoot2 = 100, LPP = 1.0, LBS) as sister to a subclade comprising all our samples of *T*. *maculipinnis*, *T*. *panchovillai*, *T*. *socolofi*, and *T*. *helleri* (Figs. 2, 3). Within this subclade all our analyses recovered *T*. *maculipinnis* as sister to (*T*. *panchovillai* (*T*. *helleri* + *T*. *socolofi*)) with high support (UFBoot2 = 100, LPP = 1.0, LBS = 100; Figs. 2, 3). All species were recovered as monophyletic, except for *T*. *helleri*, for which specimens *T_helleri_*MZ-UNICACH_1118, *T_helleri_*LSUMZ_F5718, and *T_helleri_*LSUMZ_F9509 formed a separate clade in the ML concatenated analysis, with low to moderate support (UFBoot2 = 79; Fig. 2A). This clade corresponds to *T*. cf. *helleri* in Elías et al. (2025), who also recovered these two *T. helleri* lineages based on mitochondrial cytochrome *b* data. (Fig. 2A). In contrast, in the MSC species trees (i.e., ASTRAL-IV and wASTRAL), *T. helleri* was always recovered as monophyletic and sister to *T. socolofi* (Fig. 2B). As a conservative approach for species tree inferences including one tip per species (see Methods) we grouped and named *T*. *helleri* samples according to the mitochondrial lineages defined by Elías et al. (2025; Fig. S4): *T. helleri* (*T_helleri_*MZ-UNICACH_2438 and *T_helleri_*MZ-UNICACH_2439) and *T.* cf*. helleri* (*T_helleri_*MZ-UNICACH_1118, *T_helleri_*LSUMZ_F5718, and *T_helleri_*LSUMZ_F9509) Support for relationships among *T*. *helleri*, *T*. cf. *helleri*, and *T*. *socolofi* was low across all phylogenomic inference methods (see Figs. 2, 3A). Additionally, normalized quartet scores (NQS) were similar among all possible quartets, indicating comparable frequencies of gene trees supporting alternative relationships among these taxa. The quartets recovering *T*. cf. *helleri* as sister to either *T. helleri* or *T. socolofi* had NQS values of 0.35 and 0.34, respectively, whereas the quartet recovering *T. socolofi* and *T. helleri* as sister taxa had an NQS value of 0.31 (Fig. 3A).

### Timeline of diversification of the Thorichthys-Trichromis clade

Our timetree analysis suggested that the diversification of the early-diverging Herichthyine clade *Thorichthys*-*Trichromis* took place during the Miocene epoch of the Neogene, approximately between 15 and 5 million years ago (mya; Fig. 3B). The estimated age of the most recent common ancestor (MRCA) of the two recovered *Trichromis* lineages was older (12.43 mya, during the Middle Miocene) than that of the genus *Thorichthys,* whose MRCA was estimated at 10.64 mya during the Late Miocene (Fig. 3, Table S4). Within *Thorichthys*, we estimated that both species subgroups, “meeki” and “helleri”, began diversifying around 9 mya (Figs. 3B, 4, Table S4). The ‘helleri’ subgroup accumulated its present-day diversity continuously between the Late Miocene and the Pliocene, while diversification within the ‘meeki’ subgroup (n = 3) occurred during the Late Miocene (Fig. 5).

**Fig. 4.**
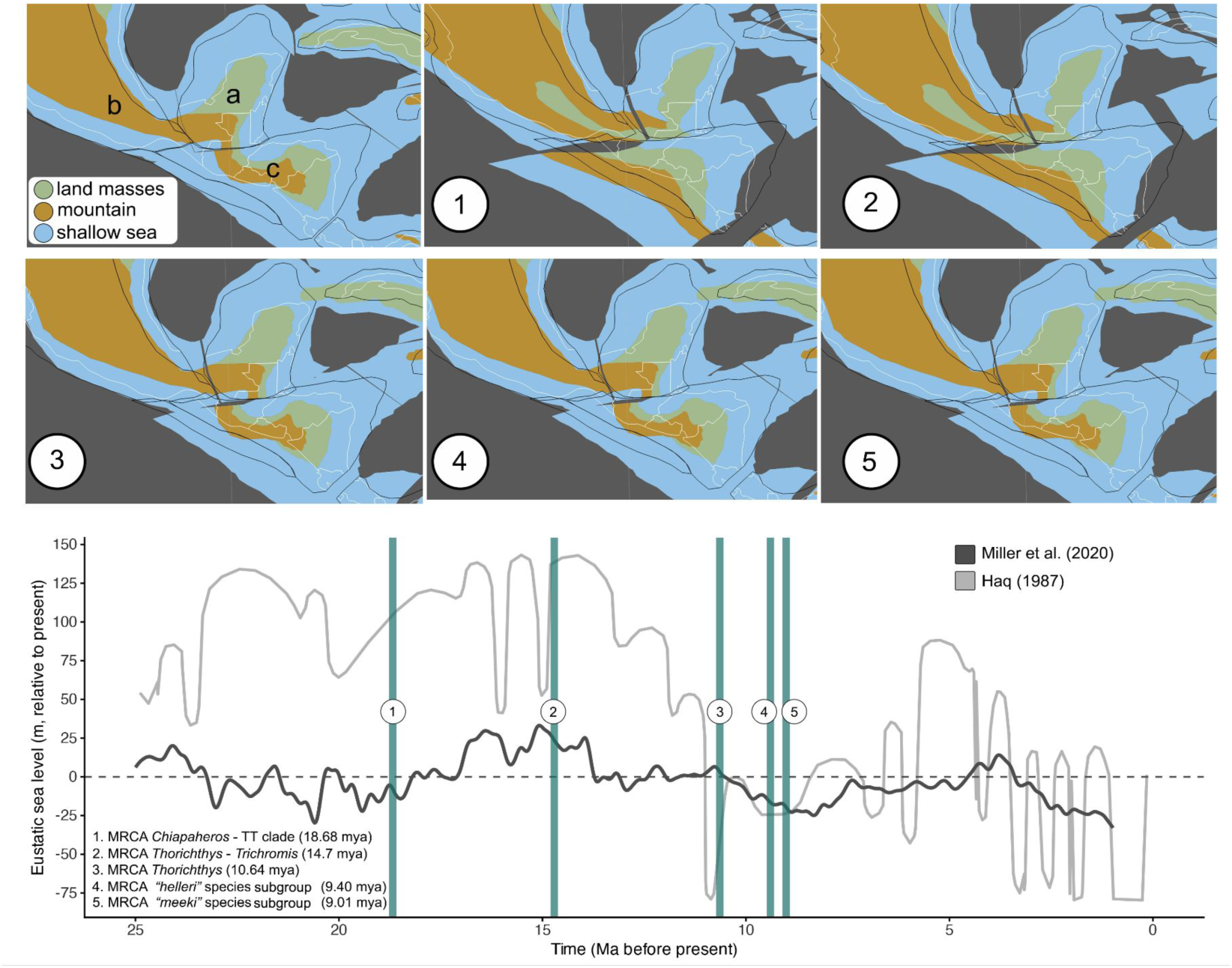
A) Paleogeographic reconstructions of the northern Neotropics based on the model of Cao et al (2017). Panels 1-5 correspond to the estimated ages of the most recent common ancestors (MRCAs) of selected nodes within the *Thorichthys*-*Trichromis*, as indicated in panel B, and show the relative position of the main geological blocks during each inferred divergence event, with map labels indicating: a, Maya Block; b, Oaxaquia-Guerrero terranes; c, Chortis block. B) Eustatic sea level changes (meters) relative to the present day (dashed line). Dark and light grey lines represent the models proposed by Miller et al. (2020) and Haq (1987), respectively, which have previously been used in biogeographic inferences of Middle American cichlids (see Discussion). Vertical green bars represent the estimated age of the MCRAs for nodes of interest of the *Thorichthys*-*Trichromis* clade and corresponding to panels 1–5 in A. Paleographic reconstructions were performed in GPlates version 2.5

**Fig. 5.**
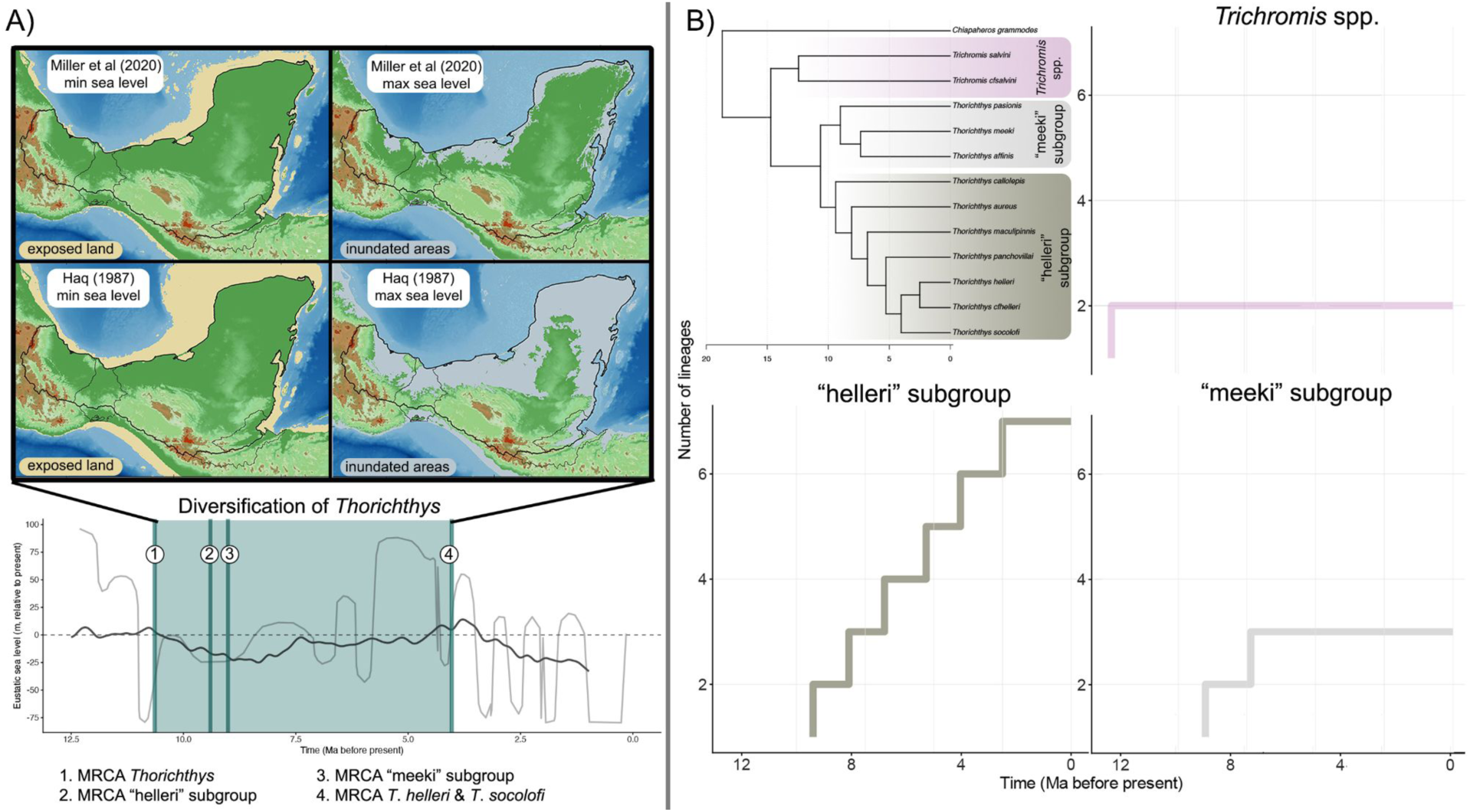
Landscape evolution of the northern Neotropics and diversification patterns of the early-diverging Herichthyines *Thorichthys*-*Trichromis* during the Middle to Late Miocene (13 to 5 million years ago). A) Hypothesized geographic landscape under two different hypotheses of eustatic sea-level changes (Haq 1987, Miller et al. 2020), highlighting exposed land and inundated areas. B) Lineage through time (LTT) plots of the early-diverging Herichthyines clade *Thorichthys*-*Trichromis*. Upper left: chronogram of the *Thorichthys*-*Trichromis* clade based on the estimated ages based on the multiple-individual Astral-IV topology (see Methods)

### Landscape dynamics in the northern Neotropics - middle Miocene through early Pliocene

The evolution of *Thorichthys* and *Trichromis* took place in a dynamic geological arena. During the earliest diversification period of the clade, during the Late and Middle Miocene, the Chortis block had not yet reached its present-day configuration (Fig. 4A). In addition, our landscape reconstructions depicted exposed continental shelves and inundated areas during the minimum and maximum sea levels estimated for the diversification period of the group. However, these reconstructions varied substantially depending on the estimates of eustatic sea-level change employed. For example, Haq, Hardenbol & Vail (1987) estimated that sea levels oscillated between +96.25 m and -79.08 m relative to present sea level between 13 and 5 mya. In contrast, Miller et al. (2020) estimated a much narrower range, between +6.9 m and -24.9 m during the same period (Fig. 5A).

### Distributional patterns of lineages within Thorichthys and Trichromis

*Thorichthys* is widely distributed across the Atlantic versant of the Maya block, with *T*. *aureus* naturally extending into the Motagua River in the Chortis block and *T*. *maculipinnis* with translocated populations north of the TMVB in the Oaxaquia-Guerrero Terrain (Figs. S1, S2). In contrast, *Trichromis salvini* does not extend to the Motagua River (Fig. S3). The two lineages of *Trichromis* are allopatrically distributed. The *T*. *salvini* ‘a’ lineage is widely distributed ranging from the Papaloapan River across the Coatzacoalcos, Tonalá, Grijalva, lower Usumacinta, the Yucatán Peninsula, and watersheds in northern and central Belize (Fig. 3B). In contrast, the *T*. *salvini* ‘b’ lineage has a relatively narrow distribution across the Lacantún, Chixoy, and La Pasión sub-basins in the upper Usumacinta and the Polochic-Cahabón AoE (Fig. 3B).

The *Thorichthys* species subgroups show contrasting geographic distributions. Species in the “meeki” subgroup (i.e., *T*. *affinis*, *T*. *meeki*, and *T*. *pasionis*) are restricted to the Grijalva-Usumacinta AoE and the Tónala River basin (Fig. S1). Within the “meeki” subgroup, *T. pasionis* is widely distributed across the Tonalá River and the western watersheds of the Grijalva-Usumacinta AoE (Fig. S1). Similarly, *T*. *meeki* is widely distributed across the Tonalá River and the entire Grijalva-Usumacinta AoE, occurring in sympatry with *T*. *pasionis* (Fig. S1), or even in syntopy. In contrast, *T*. *affinis*, the sister species of *T*. *meeki*, is restricted to the Petén lakes district and the Hondo River basin, and is allopatric to *T*. *pasionis* (Fig. S1).

The species-rich “helleri” subgroup exhibits a broader geographic distribution. Its range extends from the Papaloapan River across the Coatzacoalcos, Tonalá, the Grijalva-Usumacinta and Polochic-Cahabón areas of endemism within the Maya block, and into the Motagua River in the Chortis block (Fig. S2). Within this sub-group, the earliest-diverging species, *T*. *callolepis*, is narrowly distributed in the upper reaches of the Coatzacoalcos River (Fig. S2). Its closely related species, *T*. *aureus,* represents the easternmost distributed species of *Thorichthys*, found from the Polochic-Cahabón AoE to the Motagua River (Fig. S2). Notably, *T*. *aureus* exhibits a disjunct distribution relative to its closely related species, *T*. *callolepis* and *T*. *maculipinnis* (see Figs. 2, 3), both of which are distributed west of the Tonalá River. *Thorichthys maculipinnis*, is the westernmost distributed species, it is naturally widespread across the Papaloapan River basin with translocated populations north of the TMVB, whereas its closely related species, *T*. *panchovillai*, is widespread across the neighboring Coatzacoalcos River, east from the Papaloapan. Interestingly, *T*. *panchovillai* is sympatric with *T*. *callolepis* in the upper reaches of the basin, where syntopic populations have been reported. Finally, the clade composed of *T*. *helleri*, *T*. *socolofi*, and *T*. cf. *helleri*, is found east of the Coatzacoalcos River basin within the Tonalá and lower Grijalva rivers. *Thorichthys* cf. *helleri* has expanded its range into the Usumacinta River and riverscapes of the eastern Yucatán Peninsula (Fig. S2) where it is found in sympatry with *T*. *meeki* and *T*. *pasionis,* with documented records where all three species are syntopic.

### Ecomorphological differentiation within the Thorichthys-Trichromis clade

The principal component analysis (PCA) of four ecomorphological traits (i.e., head height, snout length, eye diameter, and body depth) revealed that 71.26% of the variance is explained by the first two principal components, with PC1 explaining 41.89% and PC2 29.38%. The remaining variance was captured by PC3 (18.66%) and PC4 (10.08%). Variation in PC1 was primarily influenced by head height and snout length (Fig. 6), with negative values correlated with taller heads and longer snouts. Whereas, PC2 variation was associated with eye diameter and body depth, where negative values corresponded with larger eyes and less elongated bodies, positive values corresponded with smaller eyes and more elongated bodies (Fig. 6).

**Fig. 6.**
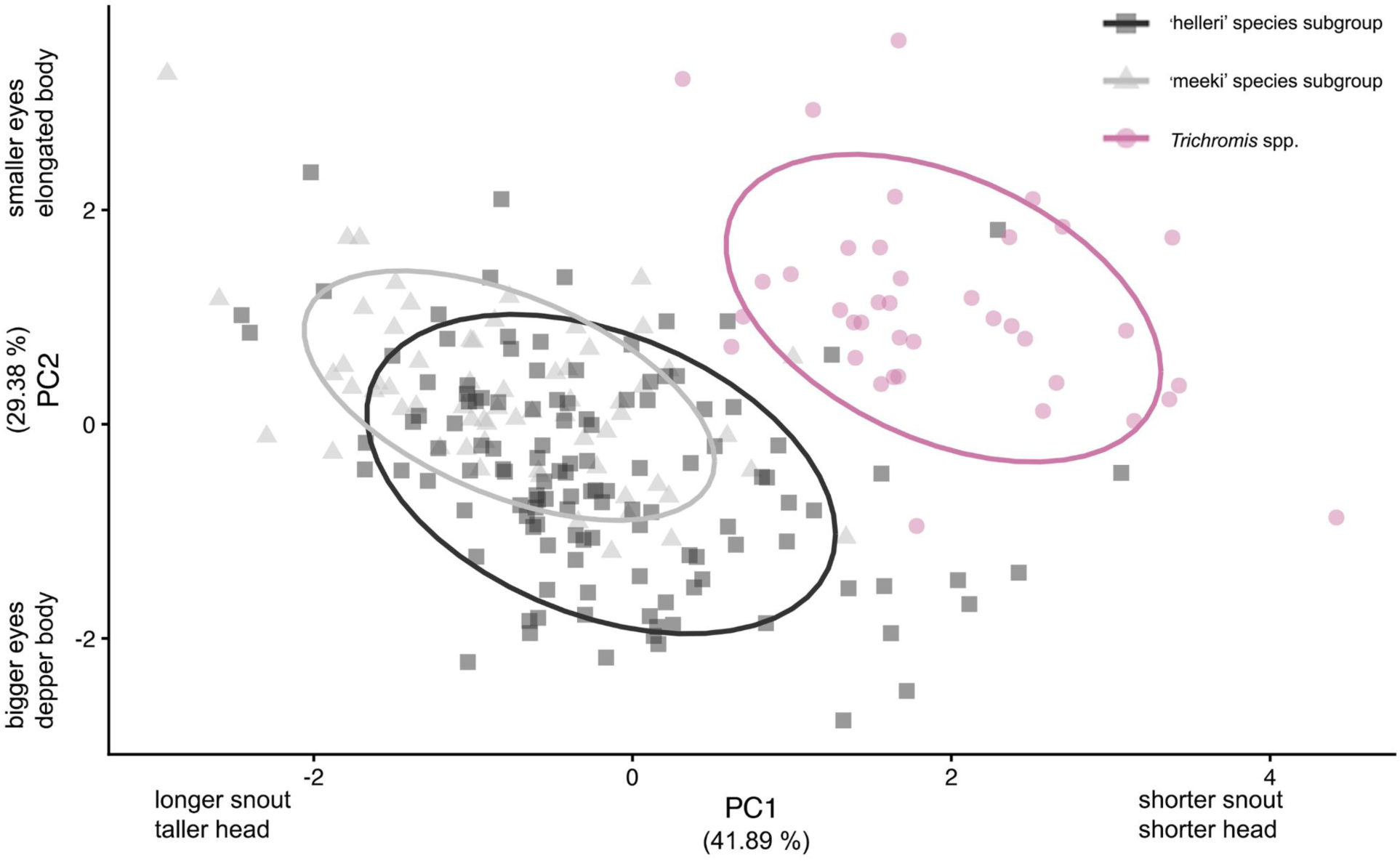
Principal component analysis (PCA) of residuals from four ecomorphological traits, snout length, eye diameter, head height, and body elongation measured from the early-diverging herichthyines *Thorichthys*-*Trichromis*. PC1 axis mainly reflects variation in snout length and head height, andPC2 primarily shows variation in eye diameter and body elongation. *Trichromis* (red circle), *Thorichthys* “helleri” subgroup (grey square), and *Thorichthys* “meeki” subgroup (light grey triangle)

The phenogram visualization of residual eye size and residual snout length showed that mean values of both traits vary among genera and between the two species subgroups of *Thorichthys*, and that both traits exhibited strong phylogenetic signal (eye size: Pagel’s λ = 1.08, p-value = 0.025; snout length: Pagel’s λ = 1.20, p-value = 4.26 x10^-5^). The two lineages of *Trichromis* possessed relatively smaller eyes and shorter snouts compared to all species of *Thorichthys* (Fig. 7). Within *Thorichthys*, species in the “helleri” subgroup had relatively larger eyes and shorter snouts than species in the “meeki” subgroup (Fig. 7). In addition, the “helleri” subgroup displayed greater variation in eye size, with several pairs of closely related species exhibiting the highest levels of divergence for this trait (e.g., *T*. *socolofi* vs. *T*. cf. *helleri*, *T*. *aureus* vs. *T*. *maculipinnis, T. callolepis* vs. *T. panchovillai*; Fig. 7A). In contrast, snout length appeared more conserved among closely related species (Fig. 7B).

**Fig. 7.**
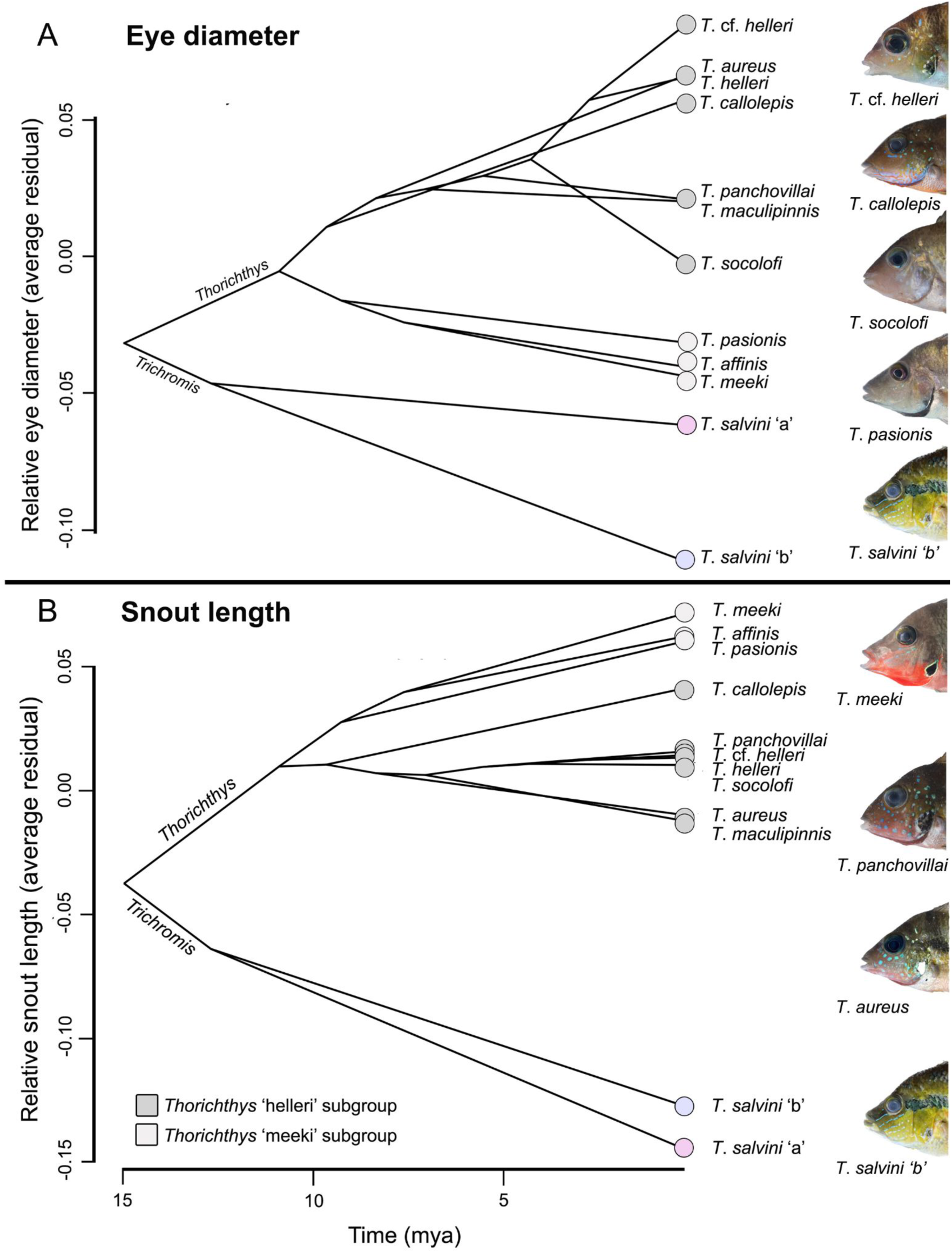
Phenograms depicting the evolution of eye diameter (A) and snout length (B) across the phylogeny of the early-diverging herichthyines *Thorichthys*-*Trichromis*. Trait values (mean residuals) are projected onto the chronogram to visualize phenotypic evolution throughout the evolutionary history of the clade

## DISCUSSION

Early-diverging herichthyine sister clades demonstrate that the diversification of northern Neotropical cichlids has been influenced by a complex interplay of introgression, extinction, ecological adaptation, geological processes, and climatic oscillations since the late Miocene. To clarify and explain these processes, we first examine patterns of phylogeny and diversity between and within the clades.

Our results support the monophyly of *Trichromis* and *Thorichthys*, revealing cryptic diversity and phylogenetic structure within both genera (Figs. 2, 3). These lineages diversified from the Late Miocene to the early Pliocene (see Figs. 3, 5), with most diversity arising west of the Usumacinta watershed. This result adds nuance to our understanding of the geographic diversification of herichthyine cichlids by emphasizing the pivotal role that river basins west of the Usumacinta (e.g., Papaloapan and Coatzacoalcos Rivers) played in their evolution. Furthermore, our results indicate that secondary contact has occurred in the upper Coatzacoalcos watershed; however, reinforcement appears to have maintained species barriers, as there is no evidence of hybridization in this region. Finally, introgression, specifically mitochondrial capture among *Thorichthys* species, likely took place within the Usumacinta watershed, potentially involving a putatively extinct lineage.

### Phylogeny and diversity of early-diverging northern Neotropical Herichthyines Genus *Trichromis*

We have inferred a robust phylogenomic hypothesis that consistently indicates that *Trichromis salvini* comprises two independently evolving lineages (Figs. 2 and 3). These findings corroborate the work of Elías et al. (2020), who identified cryptic diversity within *T*. *salvini* using mitochondrial data. By including multiple individuals (n = 13) across its distribution, we were able to broadly delineate the geographic boundaries of these two lineages, which are allopatric and exhibit contrasting distributional patterns. One lineage, *T*. *salvini* ‘a’, is widespread across most of the Atlantic versant of the Maya Block. Conversely, the lineage *T*. *salvini* ‘b’ has a narrower distribution within this area across the Upper Usumacinta and the Polochic-Cahabón biogeographic areas as described by Elías et al. (2020) (Fig. S3).

Notably, our phylogenomic hypothesis differs from those reported by Elías et al. (2020), who found mitochondrial cytochrome *b* haplotypes from the Polochic-Cahabón AoE to be more closely related to the *T*. *salvini* ‘a’ lineage, with relatively low mitochondrial divergence (cytochrome *b*: 1.07%). In contrast, our UCE-based topologies place samples from the Polochic-Cahabón and Upper Usumacinta all within the *T*. *salvini* ‘b’ clade (right-pointing and upright purple triangles, respectively, in Figs. 2 and 3), despite relatively high mitochondrial divergence between populations from these two regions (cytochrome b: 3.07%).

Mitonuclear discordance is prevalent in cichlid fishes and has been documented extensively across multiple lineages (e.g., Mbuna cichlids, Moran & Kornfield, 1993; Lamprologin cichlids, Alda et al., 2025; Jimenez et al., 2025; Heroini cichlids, Říčan et al., 2016; Mejía et al., 2025). Several non-mutually exclusive hypotheses may explain the pattern of mitonuclear discordance observed here, including cryptic diversity within the *T*. *salvini* ‘b’ lineage, with lineages allopatrically distributed between the Upper Usumacinta and Polochic-Cahabón watersheds. This hypothesis is consistent with biogeographic patterns observed in other freshwater fish genera, in which sister taxa are distributed across these two watersheds (e.g., cichlids: *Chuco*, *Cincelichthys*; poeciliids: *Carlhubbsia*, *Xiphophorus*; Rosen & Bailey, 1959; Alda et al., 2021; Elías, 2022; Du et al., 2024; Rodriguez-Machado et al., 2024). Additional explanations include differential selection on mitochondrial genomes driven by contrasting environmental conditions (e.g., temperature, dissolved oxygen) between the Upper Usumacinta and the lowlands of the Polochic-Cahabón system (Ballard & Whitlock, 2004), introgression following secondary contact, or the retention of ancestral mitochondrial polymorphism (Charlesworth, 2010) in populations across the Polochic-Cahabón.

*Trichromis salvini* was originally described by Günther (1862) based on five specimens collected by Godman and Salvin from two localities: Lake Petén Itzá and Río de Santa Isabel. Notably, these localities fall within two distinct biogeographic regions corresponding to the two lineages identified here. Lake Petén Itzá lies in the lower Grijalva-Usumacinta AoE and the distribution of the *T*. *salvini* ‘a’ lineage, whereas Río de Santa Isabel, an upper tributary of La Pasión River, one of the three basins that compose the Upper Usumacinta AoE (Elías et al., 2020), falls within the distribution of *T*. *salvini* ‘b’. *Trichromis salvini* was therefore described based on specimens belonging to two independently evolving lineages, and additional taxonomic work is warranted to resolve the identity and type locality of the nominal species. Furthermore, an integrative approach combining morphological and genomic data will be necessary to better understand and formally describe the diversity present within *Trichromis*.

### Genus Thorichthys

The two subgroups identified within *Thorichthys* differ in diversity, distribution, and morphology (Taylor & Miller, 1984; López Segovia, 2021; Elías et al., 2025). The “meeki” subgroup is relatively depauperate, comprising three species restricted to the Tonalá River and across the Grijalva-Usumacinta AoE, two of which, *T*. *meeki* and *T*. *pasionis*, are largely sympatric throughout their ranges (see Fig. S1). Interestingly, for *T*. *affinis* and *T*. *meeki,* species tree inferences (i.e., ASTRAL-IV and wASTRAL) did not support reciprocal monophyly, although the ML concatenated analysis recovered them as reciprocally monophyletic with moderate support (UFBoot2=97). This result is consistent with previous mitochondrial-based studies that failed to recover both species as independently evolving lineages and instead found low levels of genetic divergence between them (Elías et al., 2025). Given the widespread distribution of *T*. *meeki* and our limited sampling for both species (n = 8), we conservatively treat them as valid species in downstream analyses. Nevertheless, we echo Elías et al. (2025) in emphasizing that a robust species delimitation investigation is warranted to determine the number of species within the “meeki” subgroup.

In contrast to the “meeki” subgroup, the “helleri” subgroup is among the most species-rich clades of herichthyine cichlids and is broadly distributed throughout the Atlantic versant of the Maya Block, with a single species additionally occurring in the Motagua River of the Chortis block. All species within the “helleri” subgroup were recovered as monophyletic except for *T*. *helleri* (Fig. 2A). In the concatenated analysis, samples from the upper Usumacinta (*T_helleri_*LSUMZ_F5718, and *T_helleri_*LSUMZ_F9509) and the lower Grijalva (*T_helleri_*MZ-UNICACH_1118) were recovered as sister to *T*. *socolofi* rather than to *T. helleri* from the Grijalva River (Fig. 2A). In contrast, species tree analyses recovered *T. helleri* as monophyletic, although in the tree including multiple individuals per species/lineage, *T_helleri_*MZ-UNICACH_1118 from the lower Grijalva was recovered as sister to the remaining samples from the Grijalva River (i.e., *T_helleri*_MZ-UNICACH_2438, and *T_helleri*_MZ-UNICACH_2439) (see Fig. 3B). Nevertheless, the node subtending *T*. *helleri* and its sister relationship with *T. socolofi* received low support and similar NQS across all possible alternative quartets, a pattern consistent with incomplete lineage sorting which is commonly associated with recent and/or rapid speciation events (Madison & Knowles, 2006; Alda et al., 2019).

The lack of monophyly across the distribution of *T*. *helleri* has been previously reported (Říčan et al., 2016; Elías et al., 2025) and two highly divergent mitochondrial lineages (i.e., *T*. *helleri* and *T*. cf. *helleri*; ∼10% divergence) have been identified. Notably, samples showing contrasting phylogenetic placements in our analyses (*T_helleri_*LSUMZ_F5718 and *T_helleri_*MZ-UNICACH_1118) possessed cytochrome *b* haplotypes belonging to the *T.* cf*. helleri* mitochondrial lineage identified by Elías et al. (2025), whereas samples consistently recovered within *T. helleri* (MZUNICACH 2338 and MZUNICACH 2339) clustered within the nominal *T. helleri* mitochondrial clade (see Fig. S4). An ancient hybridization event between *T*. *helleri* and an extinct lineage of the “meeki” subgroup has been hypothesized to explain this deep mitochondrial divergence (see Elías et al., 2025). Based on our dating analysis (Figs. 3B), this event plausibly occurred either during the Pliocene or Pleistocene, after *T*. *helleri* expanded into the Usumacinta watershed, likely during periods of low sea-levels (Fig. 5).

To our knowledge, this represents the first report of “ghost introgression” (Zhang et al., 2019; Ottenburghs, 2020) in freshwater fishes of the northern Neotropics. Based on this evidence and the lack of support for the monophyly of *T*. *helleri,* we treated the samples that possess the *T*. cf. *helleri* mitochondrial lineage as an independent evolving lineage in our analyses. Finally, the low support and the gene-tree/species-tree discordance observed between *T*. *helleri*, *T*. *socolofi*, and *T*. cf. *helleri* (Fig. 3A) warrants further investigation into the species boundaries within these three lineages of the “helleri” subgroup distributed in the Tonalá and Grijalva-Usumacinta AoE.

Here, we propose a novel hypothesis of evolutionary relationships (Figs. 2 and 3A) for the “helleri” subgroup. Historically, the phylogenetic placement of *T*. *aureus* has remained uncertain across data types and analytical approaches. Previous molecular studies have recovered *T*. *aureus* either as sister to a clade composed of *T*. *callolepis* and *T*. *maculipinnis* (Říčan et al., 2013, 2016), or as sister to a clade comprising *T*. *helleri*, *T*. *socolofi*, and *T*. *panchovillai* (Elías et al., 2025). In contrast, morphological data have placed *T*. *aureus* as the earliest-diverging species within the subgroup (López Segovia, 2021). Our analyses instead unambiguously recover *T*. *callolepis*, restricted to the upper reaches of the Coatzacoalcos River, followed by *T. aureus*, as the earliest-diverging species within the “helleri” subgroup.

In contrast to the eastern distribution of *T*. *aureus*, its closest relatives occur in the western portion of the range for *Thorichthys*, with *T*. *callolepis* and *T*. *maculipinnis* naturally distributed in the upper Coatzacoalcos and Papaloapan rivers, respectively. We further recovered *T*. *maculipinnis* as sister to the remaining diversity of the subgroup (*T*. *panchovillai* (*T*. *helleri* + *T*. cf. *helleri* + *T*. *socolofi*))) which is distributed in the Coatzacoalcos (i.e., *T*. *panchovillai*) and across the Tonalá, Grijalva, Usumacinta and upper Sarstun rivers (i.e., *T*. *helleri* + *T*. *socolofi* + *T*. cf. *helleri*).

This novel phylogenomic pattern challenges the Usumacinta watershed as the sole “evolutionary center” for herichthyines (see Říčan et al., 2013, 2016) and instead suggests that the diversification of the “helleri” subgroup occurred outside this region, adding nuance to our understanding of the geography of diversification of northern Neotropical cichlids. Based on our dating analysis, the “helleri” subgroup began diversifying in the late Miocene to early Pliocene. Most diversification proceeded in a north-to-south direction, leading to present-day distribution of sister species across adjacent river basins, from the Papaloapan (*T*. *maculipinnis*) to the Coatzacoalcos (*T*. *panchovillai*), and further to the Tonalá, Grijalva, Usumacinta and upper Sarstun rivers (*T*. *helleri* + *T*. cf. *helleri* + *T*. *socolofi*). This temporal and biogeographic pattern coincides with periods of low sea-level during the Late Miocene (Fig. 5; Miller et al. 2020). It is therefore plausible that ancestral lineages expanded their ranges into adjacent river basins, facilitated by paleodrainage connections across the lowlands of the Maya block, followed by allopatric speciation (Dias et al., 2014; Thomaz & Knowles, 2018; Albert, Tagliacollo & Dagosta, 2020; Waters et al., 2026).

In contrast to this general pattern of diversification from the Papaloapan into the Usumacinta watershed, the biogeographic history of the early-diverging species of the subgroup (i.e., *T*. *callolepis* and *T*. *aureus*) appears more complex. In particular, the phylogenetic placement and disjunct distribution of *T*. *aureus* relative to the remaining diversity of the “helleri” subgroup is noteworthy. We propose alternative hypotheses to explain this biogeographic pattern. First, the “helleri” subgroup may have been more diverse in the past, with lineages closely related to *T*. *callolepis*, *T*. *aureus*, and *T*. *maculipinnis* potentially widely distributed across the Coatzacoalcos, Tonalá, and Grijalva-Usumacinta AoE, subsequently going extinct and resulting in the present-day disjunct distributions. Extinction has been hypothesized to be pervasive in Heroini cichlids inhabiting the lowlands of Middle America, primarily driven by marine transgressions associated with sea-level fluctuations during the early Pliocene (5.3 - 3.6 mya; see Pérez-Miranda et al., 2020). Sea level changes have been shown to influence evolutionary dynamics in freshwater fishes, promoting range expansions into adjacent river basins during low sea stands, as well as habitat loss and extinction during high sea stands (Dias et al., 2014; McMahan et al., 2017; Albert, Tagliacollo & Dagosta, 2020; Pérez-Miranda et al., 2020; Elías, McMahan & Piller, 2022; Waters et al., 2026).

Alternatively, the reconfiguration of the Chortis and Maya blocks may help explain this disjunct distribution. The geomorphological history of the Chortis block remains debated; one hypothesis suggests that it was originally positioned adjacent to present-day southern Mexico and subsequently rotated eastward to its current position since the Miocene (Keppie & Morán-Zenteno, 2005; Torres-de León et al., 2012). Given that our estimated age for the MRCA of the “helleri” subgroup falls in the Middle Miocene (9.40 mya; Figs. 3B and 4), it is plausible that the MRCA of *T*. *callolepis* and *T*. *aureus* was widely distributed across an ancestral landscape spanning present-day southern Mexico and the Chortis block. Subsequent tectonic reconfiguration, including the eastward rotation of the Chortis block, may have fragmented this ancestral distribution, resulting in a western lineage corresponding to *T*. *callolepis* (present-day upper Coatzacoalcos), and an eastern lineage corresponding to *T*. *aureus,* the most easterly distributed lineage of the genus. A similar pattern of closely related clades with wide and disjunct distributions between central Mexico and the Polochic-Cahabón AoE extending into the Chortis block, has been documented in livebearing fishes of the genus *Pseudoxiphophorus* (Agorreta et al., 2013). Rather than representing mutually exclusive explanations, these hypotheses likely reflect complementary processes operating at different temporal and geographic scales, jointly shaping the evolutionary history of the early-diverging herichthyines.

In their study of divergence times and biogeography of the herichthyine genus *Herichthys*, Pérez-Miranda et al. (2020b) concluded that cichlid divergence estimates younger or older than the bounds recovered in their study should be interpreted with caution, postulating that ages differing from their own would imply that the biogeography of Heroini cichlids is decoupled from ecology and geology. Beyond understandable criticism of data and modeling approaches, these interpretations are dismissive of alternative hypotheses and treat their proposed hypothesis as the most plausible framework to understand cichlid diversification in the northern Neotropics. Such reasoning can result in “… favouring one explanation over another simply because no other explanation was ever considered” (Parenti & Ebach, 2013). As elaborated by Parenti (2017), “biology is a powerful predictor of geology”, particularly in geologically complex regions such as the northern Neotropics. Our novel, taxonomically complete, and robust genomic dataset coupled with multifaceted paleogeographic reconstruction models corroborate a broader biogeographic pattern in which herichthyine genera have diversified outside the Usumacinta basin–including *Paraneetroplus, Vieja*, and others (Elías, 2022)–, highlighting the importance of other watersheds within the Maya block (e.g., Papaloapan, Coatzacoalcos) as evolutionary arenas for lineages within the Heroini radiation.

### Ecomorphological differentiation within the Thorichthys-Trichromis clade

*Thorichthys* and *Trichromis* occupy distinct regions of the functional morphospace (Fig. 6). Divergence in traits like eye diameter and snout length between *Thorichthys* and *Trichromis* is consistent with previous studies on functional traits of Neotropical cichlids both at macroevolutionary and ecological scales (Cochran-Biederman & Winemiller, 2013; Pease, Mendoza-Carranza & Winemiller, 2018; Soria-Barreto, Rodiles-Hérnandez & Winemiller, 2019; Arbour et al., 2020). Notably, *Trichromis* occupies a morphospace breadth comparable to that occupied by *Thorichthys*, despite having accumulated substantially lower species diversity through time (Fig. 6). Ongoing phylogeographic work combining genome-wide molecular data (e.g., ddRAD) with an expanded set of functional traits will help determine whether the morphospace occupied by *Trichromis* is geographically dispersed or clustered, thereby clarifying potential local adaptation within the genus (Zamudio, Bell & Mason, 2016).

Despite a high degree of morphospace overlap between the two *Thorichthys* species subgroups, the “meeki” subgroup occupies a narrower region than the “helleri” subgroup (Fig. 6). One plausible explanation is that species within the “meeki” subgroup have diversified under relatively similar ecological pressures, as all species are restricted to the Grijalva-Usumacinta AoE. In contrast, species of the “helleri” subgroup have a wider geographic distribution and have likely been exposed to greater environmental heterogeneity across several river basins, which may have promoted the higher morphological disparity observed.

To better understand the observed ecomorphological patterns, we focused on eye diameter and snout length, as these two traits have a large contribution to the major axes of variation (Fig. 6). Snout length exhibits strong phylogenetic conservatism, tracking both genera and species subgroups within *Thorichthys*, and shows low variation among closely related species (Fig. 7B). In contrast, eye diameter, while also showing phylogenetic signal at higher taxonomic levels (i.e., between genera and species subgroups), is the most labile trait, showing divergence among closely related lineages (e.g., *T*. *salvini* ‘a’ vs. *T*. *salvini* ‘b’; *T. socolofi* vs. *T*. *helleri* and *T*. cf. *helleri*; Fig. 7A), as well as convergence among distantly related species (e.g., *T*. *aureus* and *T*. *helleri*; Fig. 7A).

Consistent with these patterns, analyses of phylogenetic signal within the most species-rich clade, the “helleri” subgroup, revealed significant phylogenetic signal for snout length (Pagel’s λ = 1.36, p-value = 0.014), whereas eye diameter showed no detectable phylogenetic signal (Pagel’s λ ∼ 0, p-value = 1.00). Snout length, a trait commonly associated with trophic ecology in fishes (Fitzgerald et al., 2017; Lin et al., 2021), is therefore relatively conserved. In contrast, eye diameter, which is linked to habitat use and visual conditions (e.g., variation in light and turbidity regimes: Schmitz & Wainwright, 2011; Caves, Sutton & Johnsen, 2017; Andersson, Scharnweber & Eklöv, 2023), appears highly labile.

Among herichthyine cichlids, *Thorichthys* is one of the few genera in which syntopic populations have been reported. We observed that syntopic species (e.g., *T*. *callolepis* vs. *T*. *panchovillai*, and *T*. *pasionis* and *T*. *meeki* vs. *T*. cf. *helleri*) show high levels of divergence in these traits (Fig. 7). This pattern suggests that fine-scale resource partitioning (e.g., local trophic differentiation) may facilitate the coexistence of these closely related species (Begon, Mortimer & Thompson, 1996), although this hypothesis requires further ecological testing. Taken together, these results suggest that trophic and sensory traits may be shaped by different evolutionary and ecological pressures in early-diverging herichthyines.

Overall, our findings underscore that much of the diversity of northern Neotropical cichlids remains incompletely characterized, both taxonomically and evolutionarily, and warrants further exploration. This hidden diversity reflects the intricate evolutionary history shaped by the region’s complex geological landscape, which has acted as both a barrier and a conduit for diversification. The interaction of geological restructuring, climatic and sea-level oscillations, and heterogeneous ecological pressures has been pivotal in driving the evolutionary trajectories of freshwater organisms in this area. Together, these results highlight that complex geological histories and reticulate evolution jointly structure diversification patterns in freshwater systems, and emphasize the need to integrate geological and ecological complexity to fully resolve biodiversity in these systems.

## Acknowledgments

DJE thanks the E.K. Hunter Chair at Louisiana State University (LSU), the Grainger Bioinformatics Center at the Field Museum of Natural History (FMNH), and the Ohio Cichlid Association Jim Smith Fund for providing funding to conduct this research. DJE also thanks the LSU Museum of Natural Science for providing access to its molecular laboratory, and to Kevin Feldheim and Dylan Maddox in the Pritzker Molecular Lab at the FMNH. Additionally, DJE thanks the Scholarship Committee at FMNH and the Grainger Bioinformatics Center at FMNH for access to computational resources. CDM acknowledges the National Science Foundation (grant 2325892), which supported this work. All authors thank Dean Hendrickson and Adam Cohen (THNC), Benjamin Nichols and Hernan López Fernández (UMMZ), Dave Boyd and Dan Sinopoli (LSUMZ), Ben Schuster and Gretchen Hilt (SLU) who graciously assisted with loans of specimens.

## Author Contributions

**Conceptualization**: Diego J. Elías; **Data curation**: Diego J. Elías; **Formal analysis**: Diego J. Elías. **Funding acquisition**: Diego J. Elías, Prosanta Chakrabarty, Caleb D. McMahan; **Investigation**: Diego J. Elías, Sheila Rodriguez-Machado; **Methodology**: Diego J. Elías, Fernando Alda; **Data curation**: Diego J. Elías; **Resources**: Caleb D. McMahan, Isaí Betancourt-Resendes, Alejandro Díaz-Flores, Omar Domínguez-Domínguez, Ernesto Velásquez-Velásquez, Kyle R. Piller, Wilfredo A. Matamoros, Susan F. Mochel, Kevin A. Swagel, Prosanta Chakrabarty; **Visualization**: Diego J. Elías; **Writing - original draft**: Diego J. Elías, Fernando Alda; **Writing - review & editing:** Caleb D. McMahan, Isaí Betancourt-Resendes, Alejandro Díaz-Flores, Omar Domínguez-Domínguez, Ernesto Velásquez-Velásquez, Kyle R. Piller, Wilfredo A. Matamoros, Susan F. Mochel, Kevin A. Swagel, Prosanta Chakrabarty.

