## Supplementary materials for "Diversifying the Northern Neotropics: Phylogenomics and Evolutionary History of the Early-Diverging Herichthyine Cichlids *Thorichthys* and *Trichromis*"

Norte Poniente 1150, Col. Lajas Maciel, C.P. 29039, Tuxtla Gutiérrez, Chiapas, México

7 Department of Biological Sciences, Southeastern Louisiana University, Hammond, Louisiana USA

8 Instituto de Investigación en Ciencias Biológicas y Ambientales de Honduras (IBIOAH) San Pedro Sula, Universidad Nacional Autónoma de Honduras (UNAH), Honduras, 11101.

9 Museum of Natural Science, Department of Biological Sciences, Louisiana State University, Baton Rouge, Louisiana, United States of America

Running title: Evolutionary history of *Thorichthys* and *Trichromis*

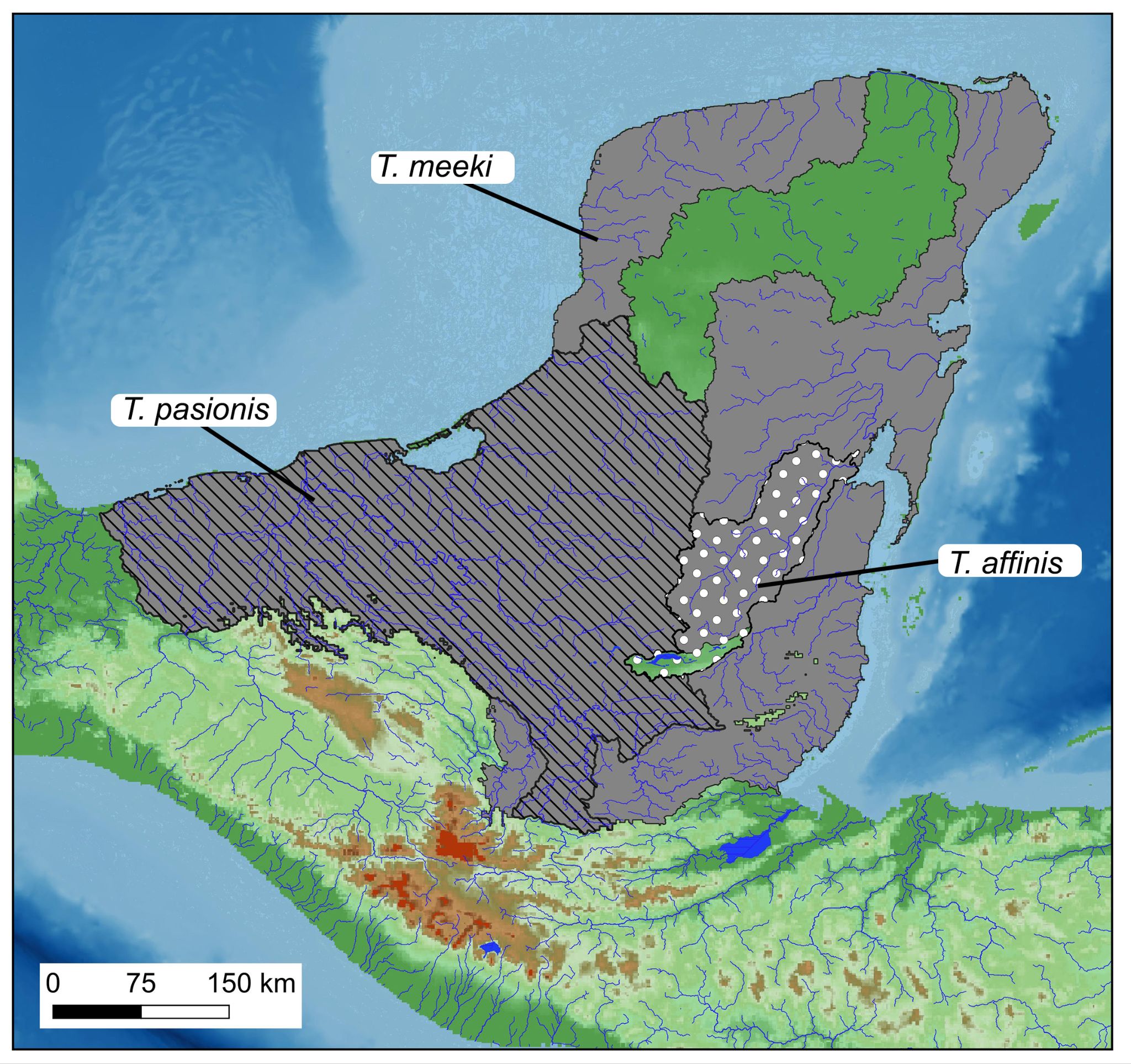

**Fig. S1** Distribution map of species of the *Thorichthys* “meeki” subgroup: *Thorichthys affinis* (white circles), *T*. *meeki* (grey shaded area), and *T*. *pasionis* (hatched area)

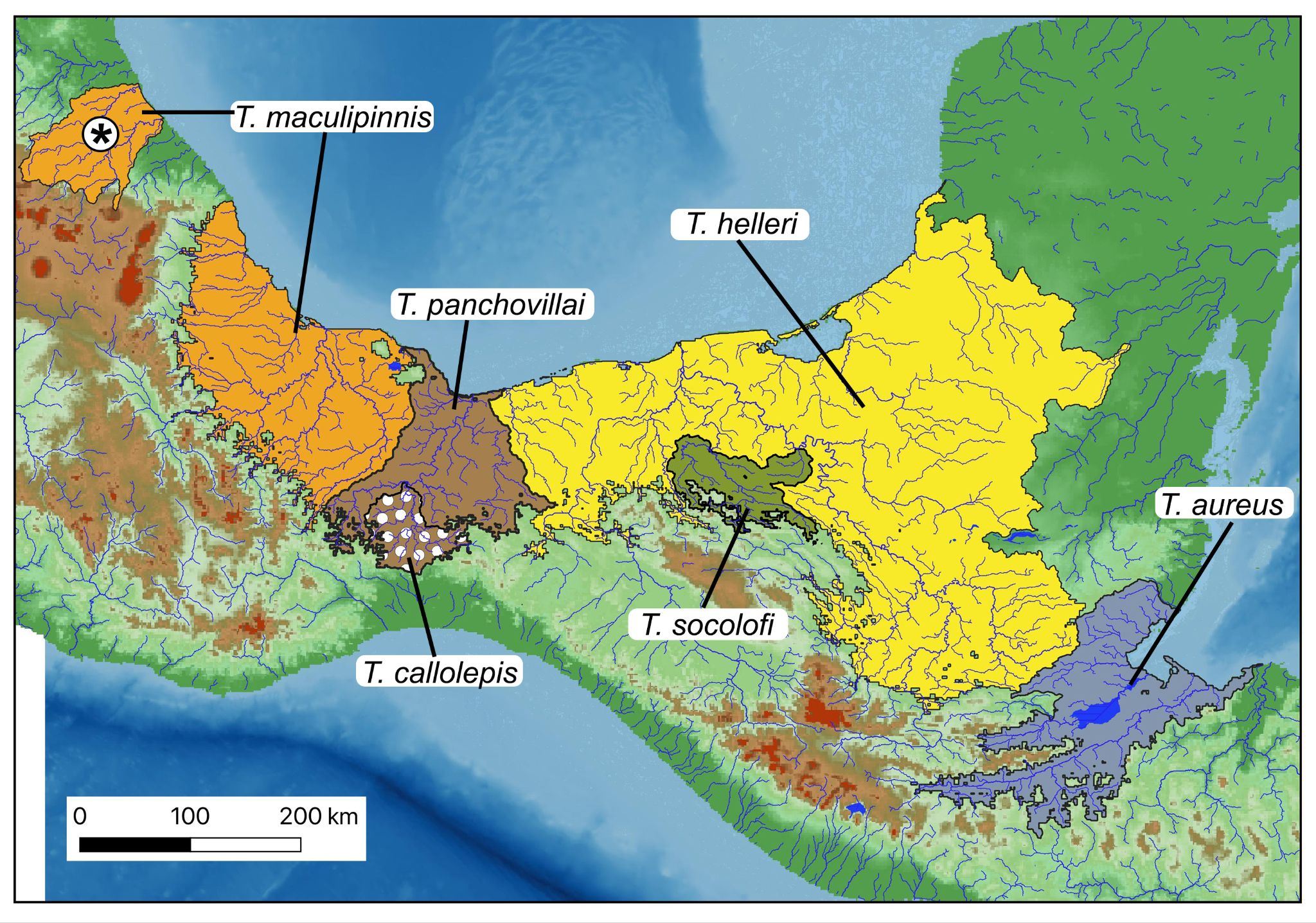

**Fig. S2** Distribution map of species of the *Thorichthys* “helleri” subgroup: *Thorichthys aureus* (blue shaded area), *T*. *callolepis* (white circles), *T*. *helleri* (yellow shaded area), *T*. *maculipinnis* (orange shaded area), *T*. panchovillai (brown shaded area), and *T*. *socolofi* (olive green shaded area). Asterisk denotes translocated populations of *T*. *maculipinnis* in Mexico, north of the Trans-Mexican Volcanic Belt.

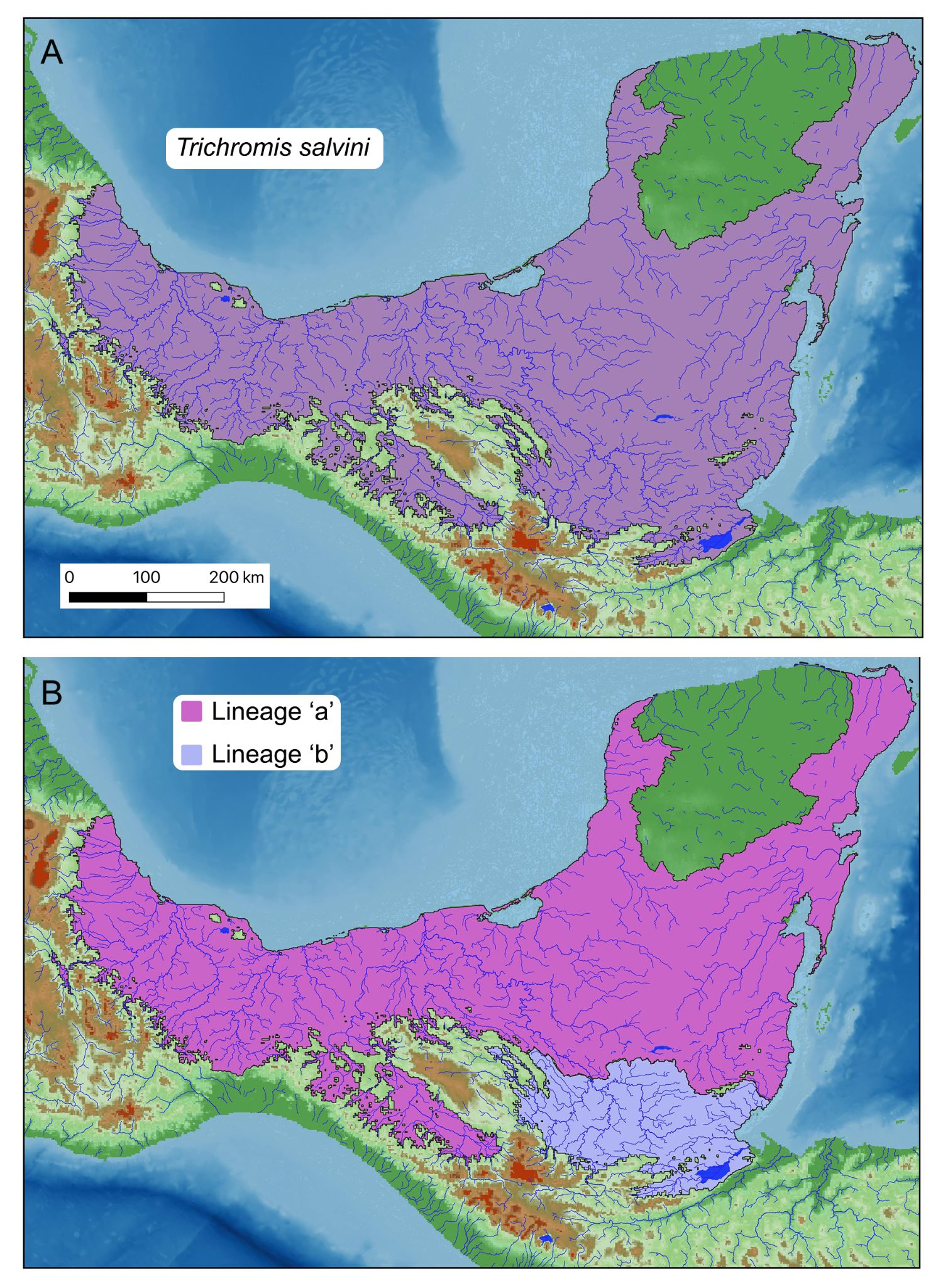

**Fig. S3** Distribution map of the genus *Trichromis*. A) Distribution of the monotypic *Trichromis salvini*. B) Distribution of the two recovered lineages within *Trichromis:* lineage ‘a’ (magenta shaded area), lineage ‘b’ (light violet shaded area)

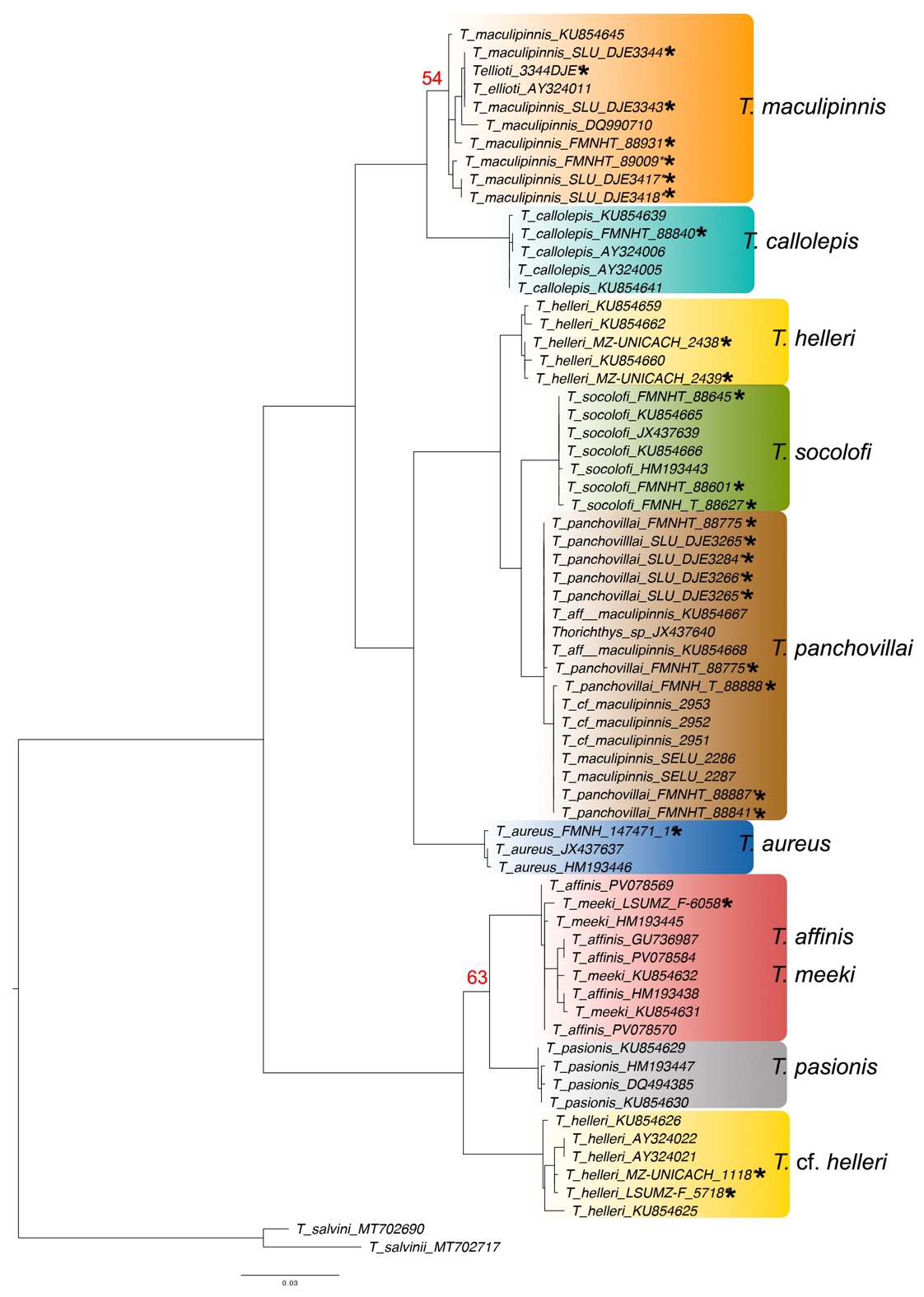

**Fig. S4**. Maximum likelihood phylogenetic inference of the genus *Thorichthys* based on novel cytochrome *b* sequences generated in this study combined with the dataset used by Elías et al. (2025). Bootstrap node support values >75 unless noted. Asterisks denote samples for which ultraconserved elements data were generated and analyzed in this study
